# Widespread dysregulation of long non-coding genes associated with fatty acid metabolism, cell division, and immune response gene networks in xenobiotic-exposed rat liver

**DOI:** 10.1101/791772

**Authors:** Kritika Karri, David J. Waxman

**Author notes:** Address correspondence to: Dr. David J. Waxman, Department of Biology, 5 Cummington Mall, Boston, MA 02215 USA. Competing Financial Interests: This work is supported in part by NIH grant ES 024421 (to DJW). The authors declare they have no other actual or potential competing financial interests.

## Abstract

Xenobiotic exposure activates or inhibits transcription of hundreds of protein-coding genes in mammalian liver, impacting many physiological processes and inducing diverse toxicological responses. Little is known about the effects of xenobiotic exposure on long noncoding RNAs (lncRNAs), many of which play critical roles in regulating gene expression. Objective: to develop a computational framework to discover liver-expressed, xenobiotic-responsive lncRNAs (xeno-lncs) with strong functional, gene regulatory potential and elucidate the impact of xenobiotic exposure on their gene regulatory networks. We analyzed 115 liver RNA-seq data sets from male rats treated with 27 individual chemicals representing seven mechanisms of action (MOAs) to assemble the long non-coding transcriptome of xenobiotic-exposed rat liver. Ortholog analysis was combined with co-expression data and causal inference methods to infer lncRNA function and deduce gene regulatory networks, including causal effects of lncRNAs on protein-coding gene expression and biological pathways. We discovered >1,400 liver-expressed xeno-lncs, many with human and/or mouse orthologs. Xenobiotics representing different MOAs were often regulated common xeno-lnc targets: 123 xeno-lncs were dysregulated by at least 10 chemicals, and 5 xeno-lncs responded to at least 20 of the 27 chemicals investigated. 81 other xeno-lncs served as MOA-selective markers of xenobiotic exposure. Xeno-lnc–protein-coding gene co-expression regulatory network analysis identified xeno-lncs closely associated with exposure-induced perturbations of hepatic fatty acid metabolism, cell division, and immune response pathways. We also identified hub and bottleneck lncRNAs, which are expected to be key regulators of gene expression in *cis* or in *trans*. This work elucidates extensive networks of xeno-lnc–protein-coding gene interactions and provides a framework for understanding the extensive transcriptome-altering actions of diverse foreign chemicals in a key responsive mammalian tissue.

## Introduction

Many industrial chemicals, drugs, environmental toxicants, and other foreign chemicals, collectively known as xenobiotics, have adverse effects on humans and other species [1, 2]. Mammalian liver is a key xenobiotic-responsive tissue: it undergoes major changes in gene expression and the epigenetic landscape, which impacts many biological pathways and induces diverse toxicological responses. Many of these responses are mediated by nuclear receptors or other transcription factors that bind foreign chemicals directly, or whose activity is indirectly altered by xenobiotic exposure [3] or by other mechanisms of action (MOAs) [4-6]. Elucidation of the gene targets, gene expression networks, and MOAs of xenobiotics is critical for the evaluation and interpretation of toxicological outcomes and hazard risk assessment.

Long noncoding RNAs (lncRNAs) comprise a significant fraction of the mammalian transcriptome and are often expressed in a highly tissue-specific or condition-dependent manner. LncRNAs act by diverse mechanisms to exert regulatory effects [7], including transcriptional, epigenetic and translational control of gene expression [8]. Many lncRNA genes are responsive to endogenous hormones [9, 10] or to environmental factors in mouse liver, as we recently observed in mice exposed to TCPOBOP, an agonist ligand of the nuclear receptor constitutive androstane receptor (CAR) [11]. It is unclear, however, whether lncRNAs respond to xenobiotics that activate receptors controlling other biological pathways, such as peroxisome proliferator-activated receptor α or γ (PPARA, PPARG), estrogen receptor (ER), and aryl hydrocarbon receptor (AhR), or to xenobiotics that act by other MOAs, including DNA damage or cytotoxicity (**Scheme 1**). LncRNA responses to chemical exposures may be MOA-specific, or alternatively, lncRNAs may respond in common to diverse chemicals that act via distinct MOAs. Further, the biological pathways associated with xenobiotic-responsive lncRNAs are almost totally unknown. Gain-of-function [12] and loss-of-function studies [13] have been used in cell culture screens to identify lncRNAs required for cell growth and drug resistance [13, 14], but such screens are not readily implemented in intact animal models. Given the widespread effects that at least some xenobiotics have on lncRNA expression [11], there is a critical need for a systematic, genome-wide approach to identify lncRNAs that respond to diverse xenobiotics in vivo, including chemicals that act via different MOAs, and to discover the biological pathways and pathological responses they may control.

Here, we use RNA-seq datasets from livers of male Sprague-Dawley rats exposed to one of 27 different xenobiotics representing seven distinct MOAs [6, 15] to discover several thousand novel liver-expressed lncRNAs and elucidate their responses to xenobiotic exposure. We identify 81 MOA-selective lncRNAs, as well as hundreds of lncRNAs that respond in common to xenobiotics acting through different MOAs. Further, we implement powerful data-driven approaches **(Figure 1)** for gene co-expression and network analysis to identify xenobiotic-responsive gene modules enriched in diverse cellular processes and to discover xenobiotic-responsive lncRNAs that occupy key regulatory points for important biological pathways commonly perturbed in xenobiotic-exposed mammalian liver.

**Figure 1.**
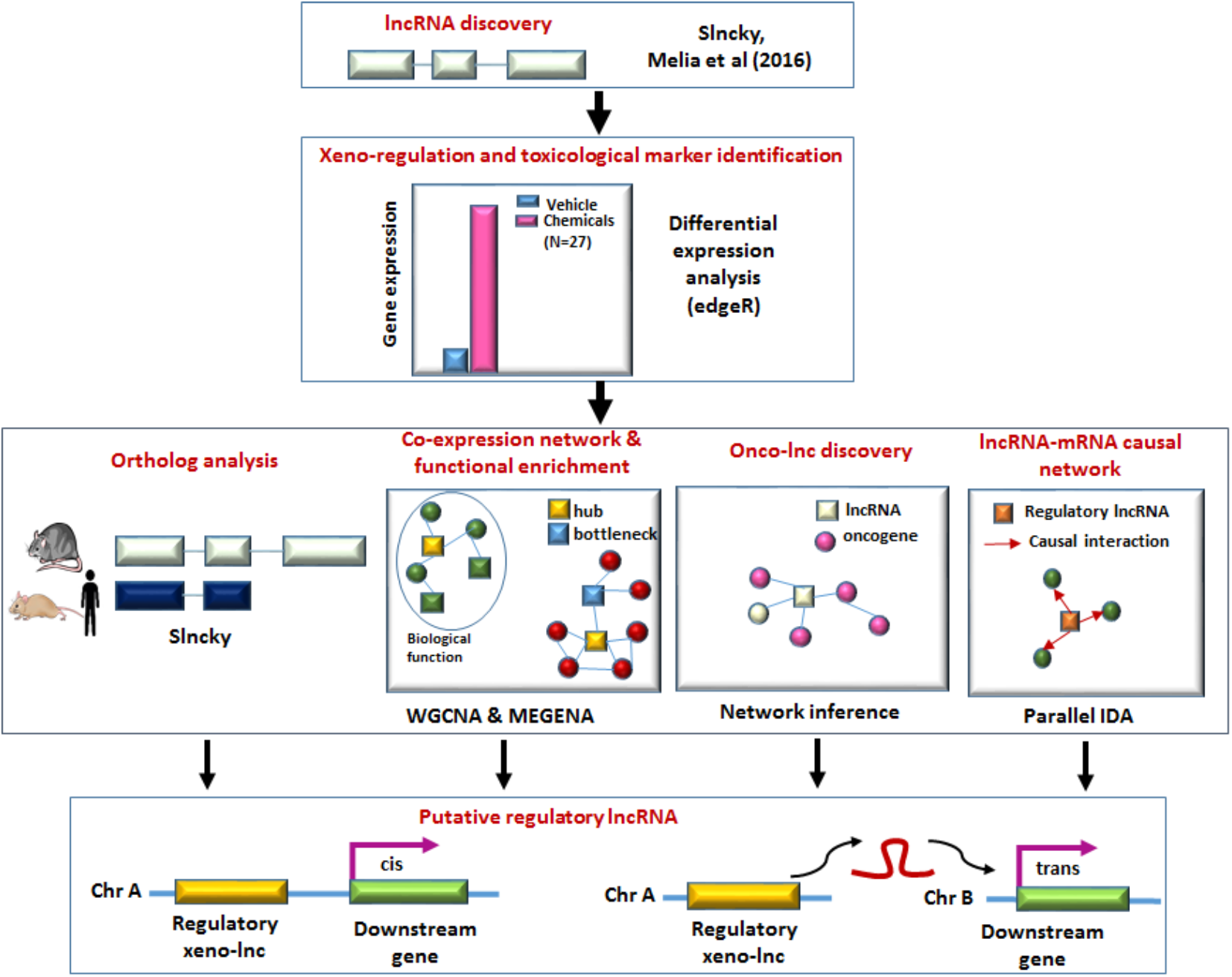
Computational framework for discovery of regulatory xeno-lncs.

## Materials and Methods

### lncRNA discovery

– 115 RNA-seq datasets from livers of male Sprague-Dawley rats exposed to one of 27 individual chemicals representing 7 MOAs (**Table S1**) were downloaded from GSE47792 [6]. Sequence reads were mapped to the rat genome (rn6) [16] using TopHat2 [17] with default parameters, including multi-mappable reads. Mapping rates ranged from 65-95% across samples. Cufflinks and Cuffmerge [18] were used to assemble a transcriptome comprised of both coding and non-coding transcripts. Transcript annotation for lncRNAs was based on three key features: transcript length >200 nt, low or no coding potential, and absence of overlap with known protein-coding genes (PCGs) [19]. Two approaches were used for lncRNAs discovery. Approach 1 [9]: First, transcripts that do not contain the above three key lncRNA features were filtered out. Thus, the 144,636 rat liver transcripts generated by Cufflinks were scanned for genes that overlap known RefSeq gene annotations. Transcripts exhibiting similarities to PCGs based on their coding potential and codon degeneracy were removed. Transcripts were converted into mature RNA sequences by exon concatenation, translated using all six open reading frames, then evaluated for the presence of protein-domain like regions (Pfam) using HMMER [20]. PhyloCSF [21] was used to identify evolutionary signatures with characteristic alignments of conserved coding regions and to distinguish PCGs from non-coding lncRNAs based on synonymous codon substitution frequencies and conservative amino acid substitutions. The coding potential of each genomic region was evaluated based on a log-likelihood ratio: transcripts with a positive score have an increased likelihood of coding for a functional protein and were discarded. The second filter excluded transcripts expressed below a fragment per kilobase length per million sequence reads (FPKM) cutoff tailored for each RNA-seq sample, which are likely to have truncated gene boundaries [9]. Genes longer than 200 nt were retained in the final set of rat liver lncRNAs. Approach 2: We used the lncRNA discovery tool Slncky [22] to identify an initial set of lncRNAs from the same 115 RNA-seq samples, filtered to remove known pseudogenes, PCGs, and artifacts from transcript assembly, and then assessed the coding potential for small peptides or novel proteins. Slncky excludes transcripts that overlap unannotated PCGs or incomplete transcripts that align to untranslated region sequences. Next, it aligns putative lncRNAs to syntenic non-coding transcripts in other, related species. Slncky then scans each significant alignment and reports back any aligned open reading frame >30 base pairs, then calculates it’s non-synonymous to synonymous substitution (dN/dS) ratio. Transcripts with dN/dS >1 have significant coding potential and are excluded. Bedtools [23] was used to determine the overlap between the lncRNAs discovered using Approach 1 and Approach 2 to obtain a combined set of 5,795 lncRNAs (**Table S2**), which includes 3,342 RNA transcripts common to both lncRNA discovery pipelines. 2,164 (37%) of the 5,795 lncRNAs are multi-exonic sequences.

### Expression quantification and clustering analysis

The full set of 115 rat liver RNA-seq samples was analyzed using a standard RNA-seq analysis pipeline. Sequence reads were mapped to the reference rat genome (rn6) using TopHat2 (v2.0.13). FeatureCounts v.1.4.6 [24] was used for read counting for both RefSeq genes (PCGs), and exon-collapsed regions of lncRNAs using a custom GTF file, available in Supplementary Materials (Rat_Liver_lncRNA.gtf). EdgeR [25] was used to identify 2,637 PCGs and 1,447 lncRNAs (xeno-lncs) that showed significant differential expression in any of the 27 chemical exposure datasets, based on these thresholds: gene up-regulation or down-regulation by at least one chemical at |fold-change (FC)| > 4, FPKM > 0.5, and Benjamini-Hochberg corrected false discovery rate (FDR) < 0.05. Differential expression data is available in **Table S3A** (xeno-lncs) and **Table S3B** (PCGs). Hierarchical clustering using Euclidean distance metric and Ward.d2 minimum variance criterion was used for clustering analysis.

### Ortholog discovery and multi-species comparisons

Primary sequence conservation was combined with syntenic conservation to discover lncRNA orthologs in the mouse and human genome, as implemented using Slncky [22]. For mouse orthologs, we considered a set of 15,558 mouse liver-expressed lncRNAs [10, 11] plus 18,065 mouse lncRNAs found in GENCODE (version M21) and 87,774 lncRNAs in NONCODE (v5). For human orthologs we considered 18,151 lncRNAs in GENCODE (version 21), 96,308 lncRNAs in NONCODE (v5) [7], and 662 human lncRNAs involved in cancer (onco-lncs) that we curated from PubMed. NONCODE (v5) [26] (http://www.noncode.org/) integrates annotations from LNCipedia [27] (https://lncipedia.org/), lncRNAWiki [28] and manual searches. Slncky uses two metrics to characterize the conservation properties of lncRNA orthologs: transcript-genome identity (TGI; percent of identical, aligning nt between a rat lncRNA and a non-transcribed locus in a syntenic region in the mouse, or human, genome); and transcript-transcript identity (TTI; percentage of identical, aligning nt present in the exonic regions of lncRNAs in both species). A TGI or TTI threshold of 30% conservation of the rat lncRNA sequence was used to define lncRNA orthologs. Putative functions for rat lncRNAs were deduced from GENCODE or NONCODE annotations [29] or from our PubMed searches for the 774 of 1,447 rat liver xeno-lncs that shared orthology with mouse lncRNAs (267 orthologs), human lncRNAs (179 orthologs), or both mouse and human lncRNAs (328 orthologs) (**Table S4**). 140 rat-mouse lncRNA ortholog pairs that respond to CAR or PXR in both species were identified as follows. First, we considered the set of 2,029 CAR/PXR-responsive rat liver lncRNAs defined at a relaxed threshold of |FC|>2 at FDR <0.05 (**Table S5A**). Of these, 381 are orthologous to one of the 15,558 mouse liver-expressed lncRNAs that we described earlier [10, 11]. Next, we identified the subset of these 381 CAR/PXR-responsive rat xeno-lncs that is found in at least one of two sets of CAR/PXR-responsive mouse liver lncRNAs, which we discovered by analyzing published RNA-seq data for expression of the above 15,558 mouse liver-expressed lncRNAs. Set 1 = 1,166 mouse liver lncRNAs that respond at |FC|>2 at FDR <0.05 to either pregnenolone 16α-carbonitrile (mouse PXR activator) or TCPOBOP (mouse CAR activator, after 3 or 27 h) [11]; and Set 2 = 1,339 mouse liver lncRNAs that respond at |FC| >2 and FDR <0.05 to TCPOBOP or to the human CAR activator CITCO after a multi-day exposure in mice at either 5 or 60 days of age (GSE98666) [30] (**Table S5B**). A total of 140 CAR/PXR-responsive rat lncRNAs had a mouse ortholog that responded to CAR or PXR activation in at least one of the mouse datasets (**Table S5C**).

### Tissue-specific co-expression networks

We used two complementary methods that employ hierarchical clustering based on gene co-expression data to assign co-regulated rat liver lncRNAs and PCGs to co-expression network modules. Weighted Gene Co-expression Network Analysis (WGCNA) [31] uses an agglomerative (bottom-up) clustering approach starting with a similarity matrix S, which is comprised of Pearson correlation values for each gene pair i and j, defined as sij= |cor(i,j)|. We transformed the similarity matrix into an unsigned adjacency matrix A by raising the correlation values to a power (‘soft’ threshold): aij = sβij with β≥1. The power β, which produces a higher similarity with a scale−free network, was used to emphasize strong correlations and punish weak correlations on an exponential scale. We selected the parameter β=12 using the pickSoftThreshold function in the WGCNA R package. Next, the adjacency matrix was converted into a distance measure (dissTOM = 1-Topological Overlap Matrix), which calculates the edge weight between two nodes based on its network neighbors and minimizes the effects of noise and spurious associations [32]. Ultimately, we identified eight rat liver xeno-gene/xeno-lnc co-expression modules (**Table S6A)** by applying the branch cutting method dynamic TreeCut [33]. WGCNA is superior in finding biological relevant modules, but has two major limitations: modules are often large, which complicates downstream refinement of biological processes; and each gene can only be assigned to a single module, which may not fully capture the complex biology, where individual genes play essential roles in more than one biological process. The second method, Multiscale Embedded Gene Co-expression Network Analysis (MEGENA) [34], addresses these limitations and generates more compact and more coherent modules, where genes can be assigned to multiple modules. MEGENA uses a divisive (top-down) approach based on the shortest path distance measure and k-medoids clustering, which finds k optimal clusters at each step by minimizing the distance within each cluster to define a more compact gene set. The clustering process continues until no further compact child clusters are formed. We used the R package MEGENA to discover 89 compact modules using the 2,637 significant PCGs and 1,447 significant xeno-lncs as input (**Figure S1, Table S6B**).

### Functional enrichment of network modules

DAVID Bioinformatics Resources 6.7 [35] was used to identify functional features of the co-expression modules discovered using WGCNA (N=8) and MEGENA (N=89). PCGs from each module were enriched for Gene Ontology terms and KEGG pathway terms (ES>1.3 and FDR <0.05) to characterize each module. Module enrichment was automated using RDAVIDWebService [36]. We labeled genes in each gene module to mark oncogenic gene drivers and tumor suppressor genes, based on a listing of 3,516 such genes that we curated from multiple sources **(Table S7**): (1) Introgen [37] collects and analyzes somatic mutations in thousands of tumor genomes to identify cancer driver genes; we identified 952 drivers with 157 druggable targets, including 689 drivers of hepatocellular carcinoma; (2) 588 tumor suppressor genes/oncogenes from Network of Cancer Genes, v6.0 [38]; (3) 1,018 genes from TSGene Database [39]; (4) 2,579 human cancer genes (allOnco; http://www.bushmanlab.org/links/genelists); (5) Cancer Gene Census catalogue, which includes genes mutations causally implicated in cancer, yielded 574 drivers or tumor suppressor genes under tier1 (i.e., gene with documented activity from experiments or literature relevant to cancers); and (6) list of 34 verified drivers and tumor suppressor genes for hepatocellular carcinoma [40].

### Hubs genes and bottlenecks ranking scheme

Biologically relevant modules based on functional enrichment results were visualized as networks using Cytoscape (v6.7) [41]. Each PCG or lncRNA is represented by a node and an edge represents a correlation value. MEGENA uses Fast Planar Filtered Network Construction [34], where significant gene correlation pairs are retained as an edge based on FDR < 0.05. For WGCNA, we selected a relatively stringent edge cutoff (|correlation| > 0.8). Topological properties of the co-expressed network modules were used to identify important functional features of lncRNAs. Nodes with a high number (high degree) of connections, called hubs, have increased likelihood of being essential [42, 43]. We also considered a second topological feature of networks, termed ‘bottlenecks’ or betweenness centrality (i.e., a measure of the number of shortest paths passing through a node), which are critical in protein networks with functional and dynamic properties control most of the information flow in a network [44]. We defined hubs as the top 10% of nodes ranked by degree, and bottlenecks as the top 10% of the nodes ranked by betweenness centrality. Cytohubba [45], a Cytoscape plugin, was used to score and rank the nodes by network features and to characterize hubs and bottlenecks for modules with functional enrichment.

### Causal Inference Network

**–** Causal inference methodology, such as Parallel IDA, outperforms methods such as statistical correlation or regression [46] and can be used to construct a lncRNA– mRNA causal network [47]. We implemented parallel IDA to infer the causal effects of xeno-lncs on target PCGs in each biological module. Parallel IDA uses intervention calculus when the directed acyclic graph is absent [48] and was used to estimate lncRNA−PCG causal relationships for co-expression modules of interest. Parallel IDA estimates the causal structure from expression data using the parallel-PC algorithm (step 1) [49, 50]; estimates of causal effects of lncRNAs on PCGs are then obtained by applying do−calculus (step 2) [51]. Step 1: We used a parallel PC algorithm and observational data (expression profiles) for the rat xeno-lncs (1,447) and xenobiotic-responsive PCGs (2,637) to learn the causal structure, which is a completed partially directed acyclic graph (CPDAG). The analysis starts with a fully connected undirected graph and then determines if an edge is to be removed from or retained in the graph by conducting conditional independence tests for the two nodes connected by the edge. We used partial correlations for the conditional independence test [52]. The algorithm then orients the CPDAG, which consist of directed and undirected edges, resulting in the equivalence class of directed acyclic graphs. The pcalg package in R was used to estimate CPDAG with a significance level (α = 0.01) for the CI test conducted by ParallelPC algorithm. Step 2: The causal effect of a lncRNA on a PCG was estimated by applying do-calculus, which when given a directed acyclic graph, estimates the causal effect of one node on any other node using observational data. Since multiple directed acyclic graphs (described above) resulted from CPDAG, we estimated causal effects for each one and used the lower bound of all possible causal effect absolute values as the final output. To construct lncRNA−PCG regulatory networks (see **Data Availability**), we used a cutoff of 0.5 for the absolute values of the causal effects to retain edges that show strong causation (**Table S8**).

### Data availability

**–** Full details for all co-expression networks and causal networks are available for visualization and query on the Network Data Exchange (NdeX) platform [53] and can be access through this link: <u>Co-expression and Causal Networks</u>.

## Results

### Liver xeno-lnc discovery

We reconstructed the transcriptome for xenobiotic-exposed rat liver based on 115 RNA-seq datasets from male Sprague-Dawley rats given one of 27 chemicals representing seven distinct MOAs (**Table S1**). We used two complementary approaches (see **Methods**) to discover a total of 5,795 liver-expressed lncRNA genes, of which 37% are multi-exonic genes (**Table S2**). 1,447 of the lncRNAs showed significant differential expression (|FC| >4 at FDR < 0.05) following exposure to one or more of the 27 chemicals, and were designated xeno-lncs (**Table S3A**). We also identified 2,637 PCGs responsive to these exposures (**Table S3B**).

### Xenobiotics grouped by MOA

We implemented hierarchical clustering of the sets of xeno-lncs and xenobiotic-responsive PCGs to elucidate gene regulatory mechanisms across the 27 chemicals. Each chemical was assigned to one of seven MOAs [6, 15] (**Scheme 1**), each represented by at least three chemicals (**Table S1**): activation of the nuclear receptors CAR and/or PXR (pregnane X receptor), considered together because of their overlapping target gene specificities [11, 54]; activation of PPAR (PPARα or PPARγ); activation of ER; activation of AhR; inhibition of HMG-CoA reductase (HMG-CoAR); non-receptor based cytotoxicity; and non-receptor-based DNA damage. Chemicals linked to PPAR, ER, and HMG-CoAR clustered tightly within their respective MOAs, whereas chemicals linked to CAR/PXR, AhR and the non-receptor based MOAs formed incohesive clusters (**Figures 2A, 2B**). Thus, the six CAR/PXR activators were separated into three separate clusters, as were the three DNA damage agents, and the AhR agonist leflunomide induced gene expression changes distinct from those induced by the two other AhR activators for both lncRNAs and PCGs. This likely reflects the diversity of mechanisms through which some of these chemicals act, e.g., leflunomide can also activate MAP kinases and induce endoplasmic reticulum stress [55]. We found 66 PCGs and 32 xeno-lncs that responded significantly to at least two of the three AhR agonists (**Table S3C**). Two of the AhR agonists, β-naphthoflavone and 3-methylcholanthrene, often showed responses opposite to those of chemicals with other MOAs, most notably PPAR activators (**Figure 2C**). Aflatoxin B1, a potent hepatotoxin and hepatocarcinogen, also has AhR agonist activity [56] and clustered more closely with those two AhR activating chemicals than did leflunomide.

**Figure 2.**
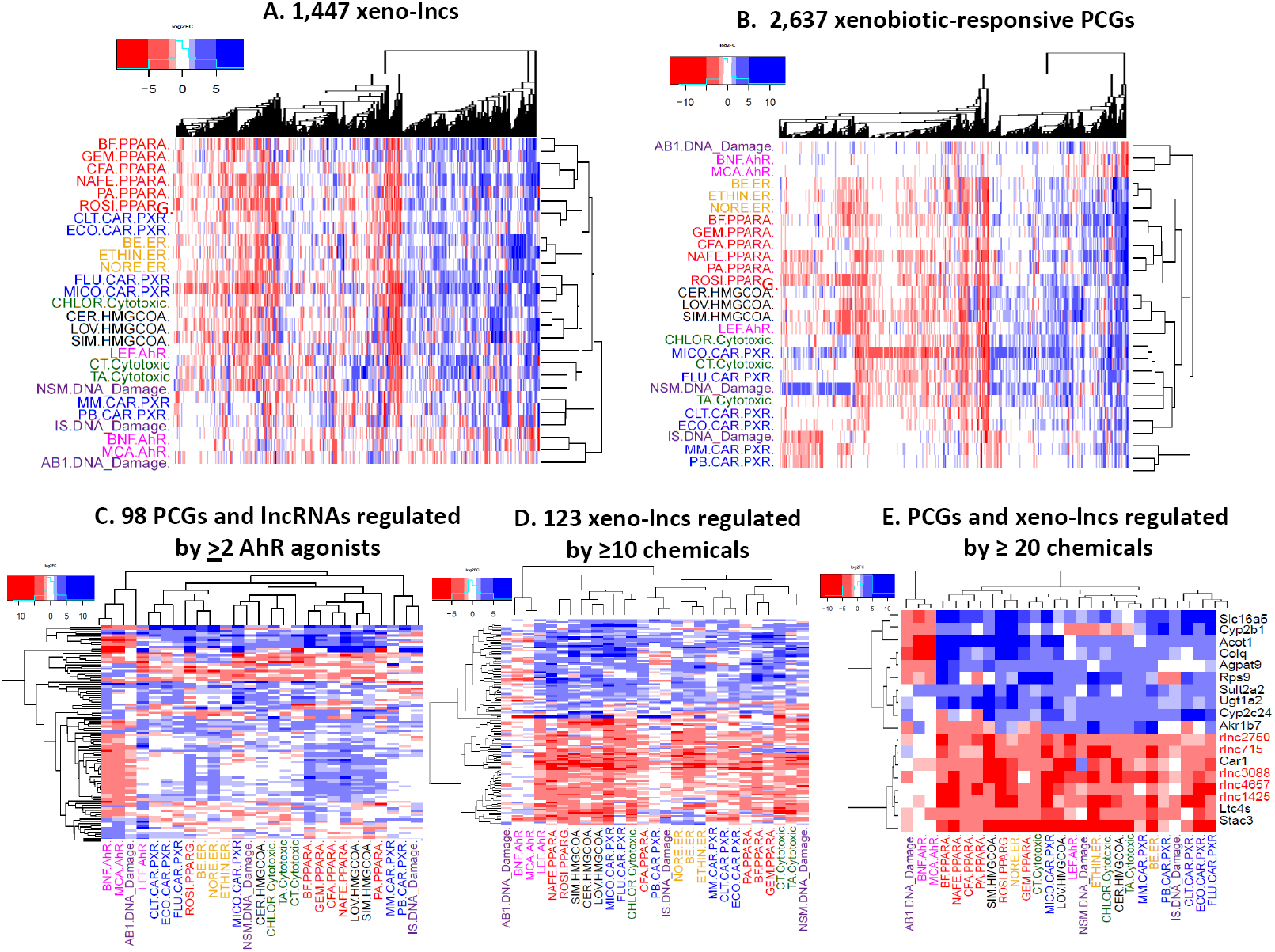
Clustering of xeno-lncs and PCGs responsive to one or more chemicals. Shown are heat maps for differentially regulated genes in rat liver that respond to 27 different chemical exposures. Chemicals are color−coded by MOA, which is indicated following the abbreviated chemical names (see **Table S1**). **A.** 1,447 lncRNAs and **B.** 2,637 PCGs up-regulated (blue) or down-regulated (red) by one or more chemicals. **C.** 32 xeno-lncs and 66 PCGs that responded significantly (|fold-change (FC)| > 4, FDR < 0.05) to at least two of the three AhR agonists. **D.** 123 xeno-lncs that respond to at least 10 of the 27 chemicals. **E.** 13 PCGs and five xeno-lncs that respond to ≥20 chemicals.

### Common xeno-lncs targets across multiple mode of actions

Many xeno-lncs were dysregulated by chemicals representing multiple MOAs. 123 xeno-lncs responded to at least 10 of the 27 chemicals tested (**Figure 2D)**. Furthermore, five xeno-lncs (rlnc4657, rlnc3088, rlnc715, rlnc1425, rlnc, and rlnc2750) were consistently down-regulated by at least 20 of the 27 chemicals (**Figure S2A)**; 13 PCGs were also dysregulated across ≥20 exposures (**Figure 2E, Table S9**). Again, gene responses to the two specific AhR agonists were similar to responses induced by aflatoxin B1 but not leflunomide. Xeno-lncs and PCGs whose expression is dysregulated by multiple chemical exposures with distinct MOAs may be associated with general responses, such as liver injury or tissue repair stimulated by xenobiotic exposure. Two of the 13 PCGs, *Sult2a2* (**Figure S2B**) and *Ugt1a2*, metabolize xenobiotics and were consistently up-regulated, while 3 other PCGs, *Ltc4s* (**Figure S2B)**, *Stac3, Car1*, were down-regulated by all but one of the 20 chemicals. The other 8 multi-xenobiotic-responsive PCGs (*Acot1* (**Figure S2B**), *Agpat9, Slc16a5, Colq, Cyp2c24*, Akr1b7, *Cyp2b1, Rsp9*) showed distinct responses to chemicals with different MOAs. Genes involved in fatty acid metabolism (*Acot1*) [57], lipid biosynthesis (Agpat9) [58], transport (*Slc16a5*), and other non-metabolic functions (*Colq*) were induced by most chemicals, but were repressed by the AhR agonists, including aflatoxin-B1. *Cyp2c24*, a rat P450, was up-regulated by chemicals with various MOAs, but showed inconsistent responses to PPAR activators (**Figure 2E**). *Akr1b7*, which detoxifies lipid peroxidation by-products [59], was induced by CAR/PXR agonists but was down-regulated by 5 of 6 PPAR activators and by four chemicals with other MOAs. *Cyp2b1* was up-regulated by nearly all activators of CAR/PXR and PPAR but was repressed by AhR agonists.

### Toxicological marker genes

We used two criteria to identify lncRNAs and PCGs that are MOA-selective markers of xenobiotic exposure: 1) the gene shows a common, robust response (|FC|>4 and FDR<0.05) to all three chemicals with the same MOA, in the case of ER, HMG-CoAR, and AhR, or to a majority (≥4) of the 6 chemicals for each of the other MOAs: CAR/PXR, PPAR, and cytotoxicity/DNA damage; and 2) the gene responds in the same direction (at a threshold of |FC|≥ 3 and FDR <0.05) to no more than 2 chemicals assigned to other MOAs, provided that the two outlier chemicals do not both act via the same MOA. We identified 162 such MOA-selective markers (81 PCGs, 81 xeno-lncs) (**Table S10**). Examples are shown for each MOA in **Figure S3**. *Ces2a* is a top marker for CAR/PXR activation, consistent with [60], while *Acox1* is a strong marker for PPAR activators, all showing >30-fold induction, consistent with findings in mouse and human models [61]. Further, two xeno-lncs were markers for both CAR/PXR and PPAR. *Aldh3a1* was a marker AhR responses, consistent with its strong induction by hepatotoxic dioxin-like compounds [62]. Finally, *Mapk15* was strongly induced by all three chemicals that target HMG-CoAR, *Myom2* was a marker for ER agonists, and *CD300lb* was a marker for chemicals that induce cytotoxicity or DNA damage. These marker genes may be incorporated into toxicological signatures to characterize MOAs of novel xenobiotic exposures.

### Identifying functional xeno-lncs by evolutionary analysis

Ortholog discovery and comparative studies across species are widely used for functional annotation of evolutionarily conserved PCGs. However, lncRNAs evolve rapidly, and many are not broadly conserved [63]. LncRNAs are, however, often found in syntenic positions in the genome and share short conserved domains [64]. Here, we used a combination of sequence identity and synteny between species to identify orthologs of the rat xeno-lncs and then infer biological functions. Of 5,795 liver-expressed rat lncRNAs, 3,020 (52%) shared orthology with either mouse or human lncRNAs, including 774 (53%) of the 1,447 rat xeno-lncs (**Figure 3A**). Some of these orthologs are well-characterized functional lncRNAs. For instance, rat xeno-lnc rlnc449, which is induced by HMG-CoAR inhibitors and cytotoxicity and DNA damage agents but is repressed by several activators of CAR/PXR, PPAR and ER, showed 64% sequence identity with human lncRNA H19, both at the transcript level (TTI) and at the genome level (TGI) (**Figure 3B**). LncRNA H19 is important for fetal liver development and is repressed after birth, but is over-expressed in multiple human cancers, including hepatocellular carcinoma, where it enhances tumor progression [65, 66]. Another rat liver xeno-lnc, rlnc397, which is induced by several nuclear receptors agonists and by the hepatoxin carbon tetrachloride, shares 66% TTI with mouse lncRNA Snhg14 (**Figure 3C**). Snhg14 is overexpressed in several cancers and potentiates tumor progression, in humans, by serving as a sponge (endogenous competitor RNA) for multiple microRNAs [67-69]. **Table S4** presents the full set of rat lncRNA orthologs and their inferred functions.

**Figure 3.**
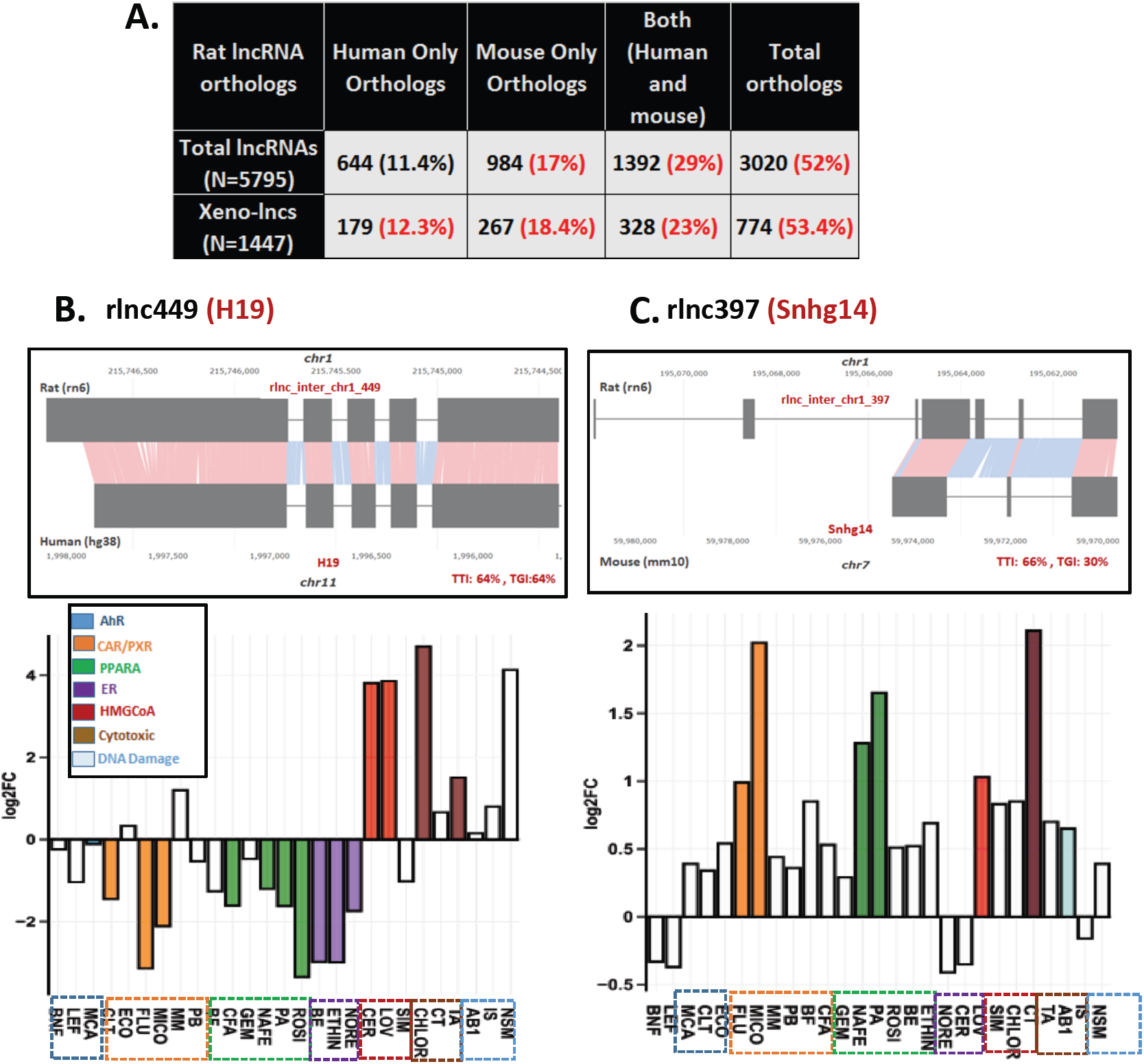
lncRNA orthology. **A.** Shown are the number of mouse and human orthologs discovered for 5,975 rat liver lncRNAs and the subset of 1,447 xeno-lncs, based on a TTI or TGI cutoff >30%. **B.** Rat rlnc449 is orthologous with human lncRNA H19. Shown is the alignment between the two orthologs (top) and rat liver expression data for rlnc449 across 27 chemicals (bottom). Colors are used to mark expression data for chemicals in each MOA. Gene responses (log2 FC values) are shown on the y-axis. White bars represent non-significant responses (FDR≥ 0.05). **C.** Rat rlnc397 aligned with its mouse ortholog lncRNA Snhg14, and expression data for rat liver rlnc397, as in B.

### Rat-Mouse ortholog responses to CAR/PXR activators

We identified 140 rat xeno-lncs with mouse orthologs that responded to at least one CAR/PXR activator in both mouse and rat liver. Two−way hierarchical clustering of expression data across exposures indicates species is a dominant separation factor (**Figure S4A**). Some lncRNAs showed a consistent pattern of dysregulation in both species (rlnc4048−mlnc4655, both up (**Figure S4B**); and rlnc4100−mlnc4577, both down (**Figure S4C**)), while others showed opposite regulation (rlnc2209-mlnc3859; **Figure S4D**). Some xeno-lncs were responsive to CAR activators after multiple days of exposure, e.g., rlnc1448−mlnc2065, which responded to CAR activators significantly (|FC| >2, FDR <0.05) in both species after a 4-5-day exposure, but not after 1 day in mouse liver (**Figure S4E**, Table S5C). Thus, xeno-lncs perturbed by chemicals with the same MOA can exhibit different responses, depending on species and duration of exposure.

### Functional prediction of lncRNA−PCG relationships using gene co-expression networks

Functional lncRNAs have been identified using cell-based screens, e.g., for effects on cell growth, however, that approach is not readily implemented in an intact liver model. Here, in an alternative approach, we clustered the 1,447 rat xeno-lncs together with the 2,637 xenobiotic-responsive rat PCGs based on their expression profiles across the 27 chemical exposures. This allowed us to infer rat xeno-lnc functions from the known functions of PCGs in the same co-expression cluster (guilt-by-association) [70, 71]. We constructed co-expression networks using two complementary methods: WGCNA, which uses agglomerative (bottom-up) clustering, resulting in large gene modules [31]; and MEGENA, which uses divisive (top-down) clustering to discover smaller, coherent network modules and sub-modules [34]. We identified eight co-expression modules (gene networks) using WGCNA and 89 using MEGENA (**Table S6A, S6B**). For each co-expression network, we identified rat lncRNAs whose orthologs have functional annotations, including oncogenic gene drivers and tumor suppressors. We also computed module functional enrichments to obtain a primary level of annotation for lncRNA function. These lncRNA−PCG networks helped us identify many highly connected lncRNAs in each module, including intra-modular hubs and bottleneck genes, which are at critical nodes controlling communication among other nodes in the network, as indicated by a high number of non−redundant shortest paths through the specific node or edge [44]. We discovered co-expressed modules with striking functional enrichments for genes in multiple biological processes and pathways, including fatty acid metabolism (MEGENA module C9), lipid and sterol metabolism (module Black), cell cycle (module C13) and immune response (module C7) (**Table S11, Table S12**). One module, C10 (68 PCGs, 91 xeno-lncs), did not show any significant biological or pathway enrichment, but contains all 50 ER marker genes. The eleven xeno-lncs identified as hub or bottleneck genes in this module are suggested to play a regulatory role in estrogen-based pathways (**Figure S5, Table S12A**). Highlights of select gene modules are presented below.

### Xeno-lnc hub and bottleneck genes regulating fatty acid metabolism and PPAR signaling

Module C9 (258 PCGs, 152 xeno-lncs) was strongly enriched for genes involved in fatty acid metabolism (N=114 genes, Enrichment Score (ES) =18.5, **Table S11E**). This module is also strongly enriched for peroxisomal genes (ES=13.0) and harbors 35 of the 39 (90%) PPAR marker genes. Seven xeno-lncs were either top intra-modular hubs, bottlenecks, or both (**Figure 4, Figure S6, Table S12B**), which indicates they have important regulatory roles and their xenobiotic responsiveness may contribute to dysregulation of hepatic fatty acid metabolism following chemical exposure [72]. One of these lncRNAs, rlnc973, is orthologous with human lncRNA AS-RBM1, which enhances RBM1 protein translation and up-regulates megakaryocyte differentiation and leukemogenesis [73]. 35 xenobiotic-responsive PCGs in this module are also critical hub or bottleneck genes involved in fatty acid metabolism (**Table S12B)**, and five of these genes (*Cyp2j4, Acat1, Acaa1a, Ucp3, Acox1*) are PPAR MOA-selective marker genes (**Table S10**).

**Figure 4.**
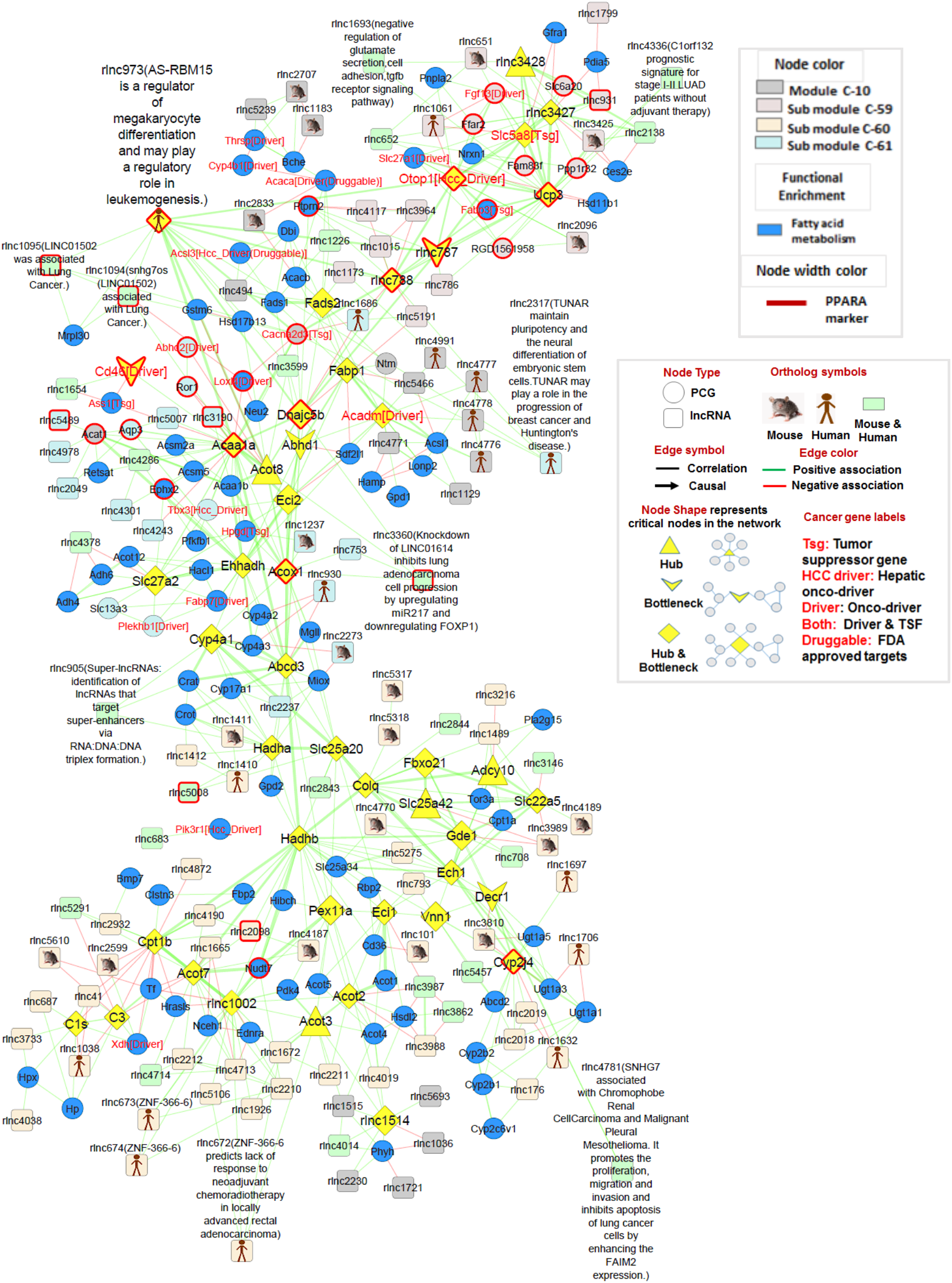
Fatty acid metabolism-enriched module C9. Fatty acid metabolism genes are marked with blue nodes. Network submodules (C−59, C−60, C−61) are represented by other colors, as indicated. Thirty−five nodes in the network are PPAR marker genes (dark red node border color). lncRNA hubs or bottlenecks: Three xeno-lncs in the module are hubs (rlnc3428, rlnc788, rlnc787) and four are both hubs and bottlenecks (rlnc3427, rlnc1514, rlnc973, rlnc1002). The full network is shown in Figure S6; this excepted segment shows xeno-lncs directly connected to PCGs that are either: involved in fatty acid metabolism, PPAR markers, or are hub or bottlenecks. The legend (in box) describes nodes and edge properties in co-expression and causal networks. Nodes represent genes or xeno-lncs, and an edge shows the relationship between two nodes, with a correlation or absolute value of causal effect (causation). PCGs and xeno-lncs are represented as network nodes with circle and square shapes, respectively. LncRNA orthologs (human, mouse, or both human and mouse) are marked using different symbols. Edges in the co-expression network correspond to correlation values between the two connected nodes (genes or lncRNAs), marked as a line. Arrows are used in causal networks to indicate causality between regulatory lncRNAs and their PCGs targets. Green edges show positive associations and red edges represent negative associations. Critical nodes (hubs, bottlenecks, or hub−bottlenecks) are shown using different node shapes and with a node color. PCGs that are either cancer drivers or tumor suppressors are labelled in red text enclosed in square brackets. The same legend applies to Figures 5-8.

### Xeno-lncs associated with cell cycle regulatory genes

Module C13, comprised of 106 PCGs and 6 xeno-lncs, includes 87 PCGs (82%) involved in cell cycle pathways (**Figure 5A, Table S12C**). The strong enrichment of this module for cell cycle and cell division genes (ES= 17.7) and for microtubule binding, spindle, kinesin and related terms (ES=7.6) (**Table S11H**) implicates the six lncRNAs in this module in these processes. One, xeno-lnc rlnc3347, is orthologous with NORAD, a human lncRNA that is up-regulated by DNA damage, maintains chromosomal stability in human cells, and is implicated in tumorigenesis [74, 75]. 33 of the 87 cell cycle regulatory genes in module C13 are oncogenic drivers and two are tumor suppressors (**Table S13**). Five of the six xeno-lncs may be oncogenic (onco-lncs), as they show significant positive correlation with cancer drivers in the subnetwork, albeit weakly (**Figure S7**). Down regulation of the oncogenic drivers and the onco-lncs in module C13 was commonly seen after exposure to chemicals associated with HMG-CoAR, ER, AhR, PPAR and CAR/PXR (**Figure 5B**). One exception was rlnc3587, which was significantly up-regulated by 10 of the chemical exposures (**Figure 5B**, red arrow). All 35 oncogenic drivers and tumor suppressors were strongly induced by the hepatocarcinogen N-nitrosodimethylamine (log_2_|FC| = 2−5), and to a lesser extent by PPAR activators and CAR/PXR-activating chemicals. The finding of multi-xenobiotic responsive xeno-lncs in this module highlights the potential cell cycle disruptive actions of the associated xenobiotics.

**Figure 5.**
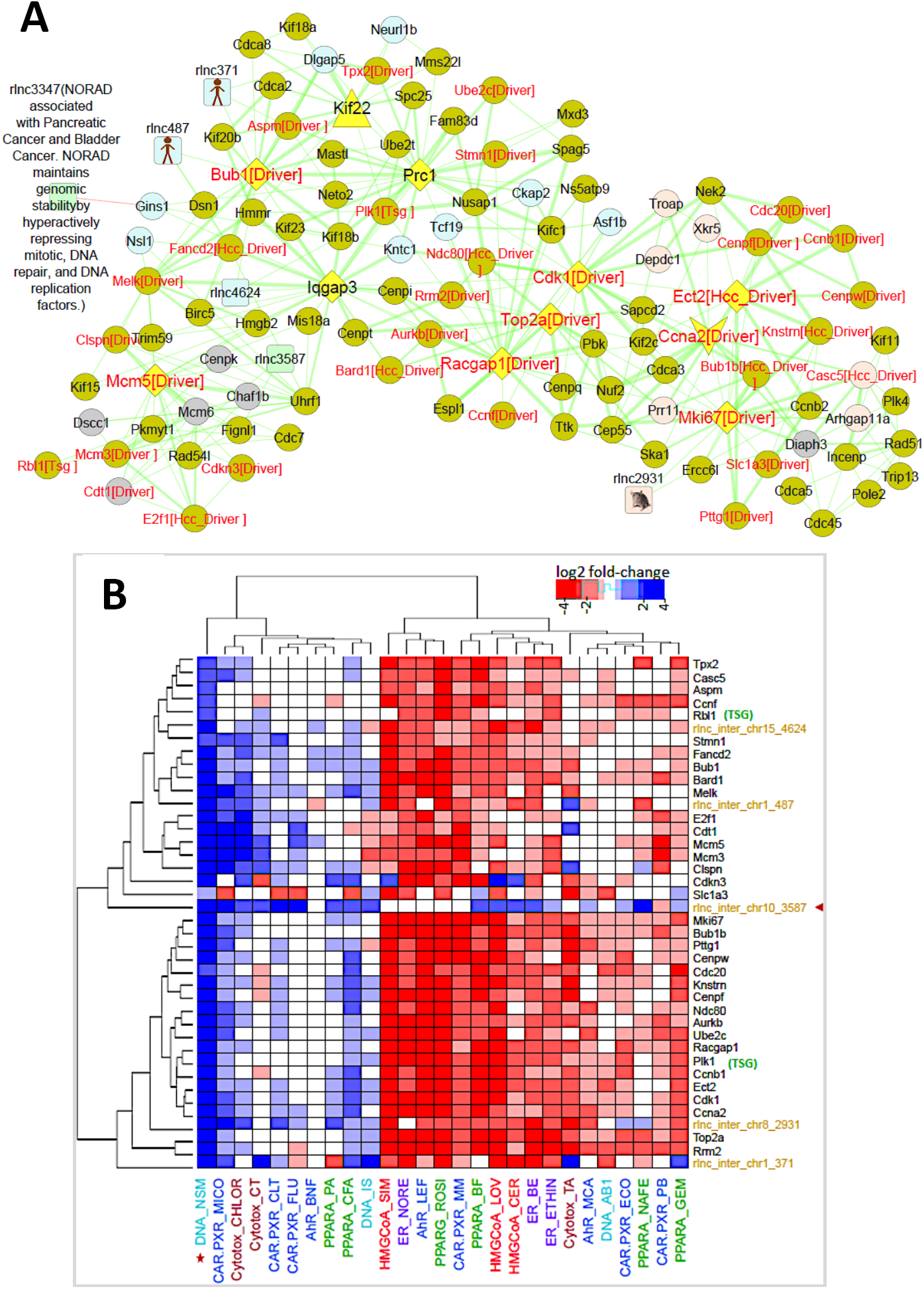
Cell cycle-enriched module C13. **A.** Shown is module C13 (PCG: 104, xeno-lncs: 6). **B.** Heat map showing two-way hierarchical clustering of gene responses (in log_2_ FC) for oncogenes and their connected xeno-lncs from module C13, across 27 chemical exposures. Xeno-lnc rlnc3587 (red arrow) was up-regulated by 10 of the chemicals.

### Immune based responses of xeno-lncs

Module C7 (480 PCGs, 94 xeno-lncs) was highly enriched (ES=14.2) for immune response and related terms (chemokine, cytokine and MHC processing) (**Table S11D, Table S12D**). This module includes 94 xeno-lncs, of which three are hub genes (rlnc2830, rlnc1130, rlnc1023). rlnc2830 was positively co-expressed with nine PCGs involved in immune response (**Figure 6A**). These include factors active in lymphocyte signaling and antigen uptake (Hcls1) [76], regulatory T cell-mediated suppression of CD4+ effector T cells (Ncf1) [77], modulation of macrophage functions (Slamf8) [78], and T cell-dependent immunity (Hk3) [79]. N-nitrosodimethylamine, which activates lymphocytes to produce pro-inflammatory cytokines [80] that induce hepatic fibrosis and liver inflammation [81], induced rlnc2830 and all nine immune genes (**Figure 6B, Table S14A**). N-nitrosodimethylamine also up-regulated 112 (64%) of the 125 oncogenic driver or suppressor genes in module C7 (**Table S14B**). The hub gene rlnc1130 was connected to six genes in module C7, while hub−bottleneck rlnc1023 made connections with a partially overlapping set of nine genes **(Figure 6C)**. rlnc1023 negatively correlated with Arg1, an immunosuppressive gene [82], and with Emp2, a tumor suppressor (<u>Li et al. 2013</u>). These two regulatory xeno-lncs were negatively correlated with Sox4, a heptaocarcinogenic driver [83], and showed positive associations with Il6R and Mat1a. Miconazole, an antifungal agent and CAR/PXR agonist, significantly repressed both regulatory xeno-lncs and seven of the PCGs, while strongly inducing three others, including S100a6 (**Figure 6D**). Increased expression of S100a6 promotes cell proliferation in human HCC [84]. Overall, 50 xeno-lncs from module C7 showed either positive or negative associations with 125 oncogenic drivers or suppressors (**Table S14B**). Chemicals that dysregulate these xeno-lncs are expected to have a major impact on tumorigenesis.

**Figure 6.**
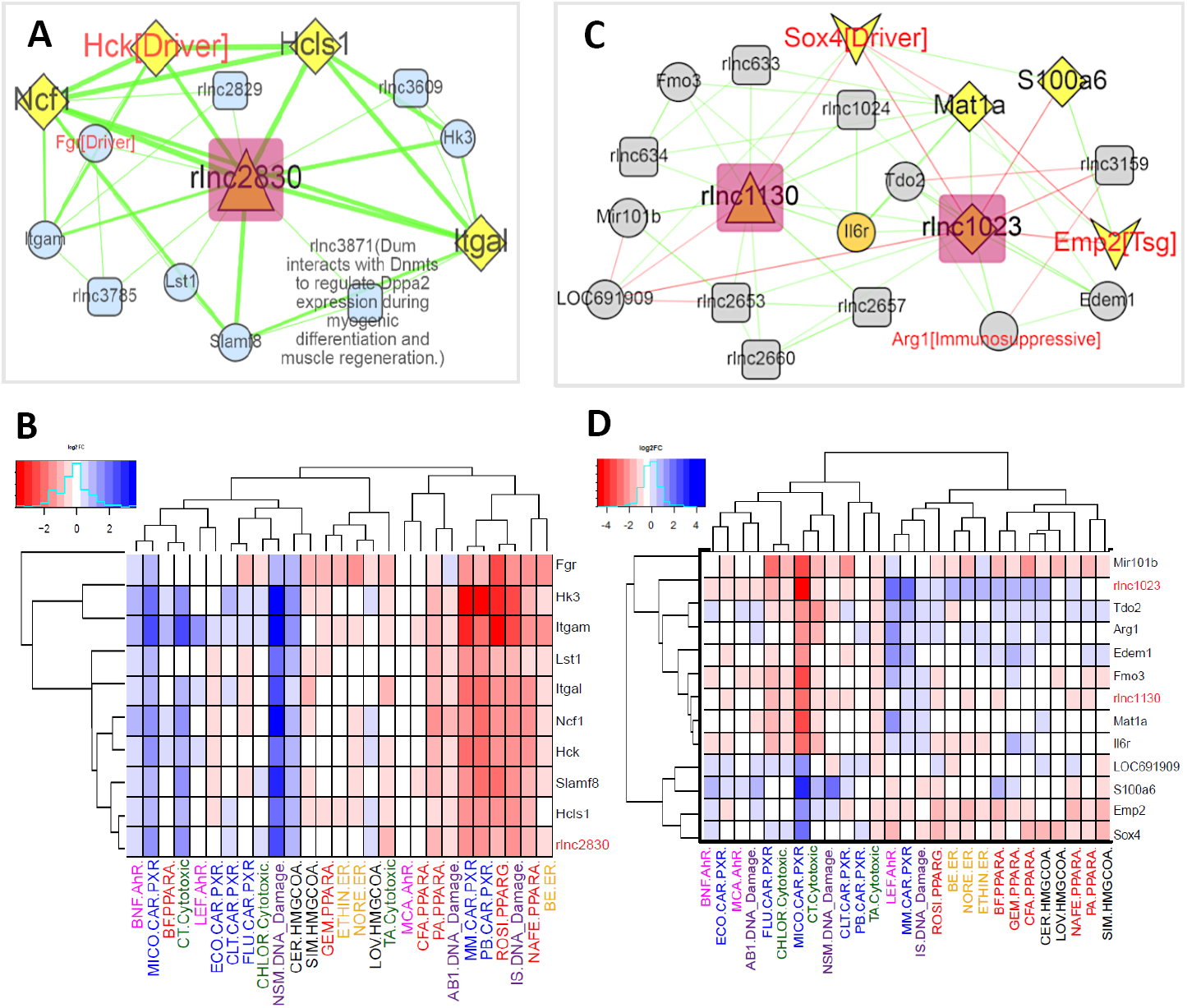
Subnetworks involving hub and bottleneck genes in module C7. **A.** rlnc2830, a hub gene from module C7, is positively co-expressed with nine PCGs involved in immune response. **B.** Responses of rlnc2830 and its PCG partners across 27 chemicals (in log2 FC). **C.** rlnc1130, a hub gene connected to six genes in module C7 and rlnc1023, a hub−bottleneck gene with connections to nine genes. **D.** Responses of rlnc1130 and rlnc1023 and their PCG partners across 27 chemicals (in log2 FC).

### Regulatory xeno-lncs associated with sterol metabolism and viral response

We used a parallel IDA algorithm to learn causal structures from expression data and construct lncRNA−PCG causal regulatory networks for biologically interesting modules. Xeno-lncs that occupy critical nodes in co-expression modules (i.e., hubs or bottlenecks) often showed strong causal effects. For example, WGCNA module Black contained 145 nodes (103 PCGs, 42 xeno-lncs). Causal network analysis based on an absolute value of causal effect cutoff >0.5 reduced the Black module to 90 nodes, including 25 xeno-lncs showing strong causal effects on their predicted gene targets (N=65). This module was enriched in sterol metabolism (ES=13.7, **Table S11K**; pink nodes in **Figure 7**) and encompassed 36 of 63 (57%) HMG-CoAR marker genes (black node border color, **Figure 7**; **Table S12E**). A small isolated sub-network was related to viral response (ES=4.1; green nodes), with rlnc4746 acting as the causal center. Two regulatory xeno-lncs in the causal network (rlnc2973, rlnc322) have functional orthologs positively associated with tumorigenesis: Lnc-SC5DL-3:1 (ortholog of rlnc2973) is up-regulated in triple negative breast cancer [85]; and elevated expression of lncRNA RP11*-*21L23.2 (ortholog of rlnc322), is associated with high risk in non-small cell lung cancer [86]. Both xeno-lncs were strongly induced (log_2_ FC = 4−6) by all three HMG-CoAR inhibitors (**Table S3A**).

**Figure 7.**
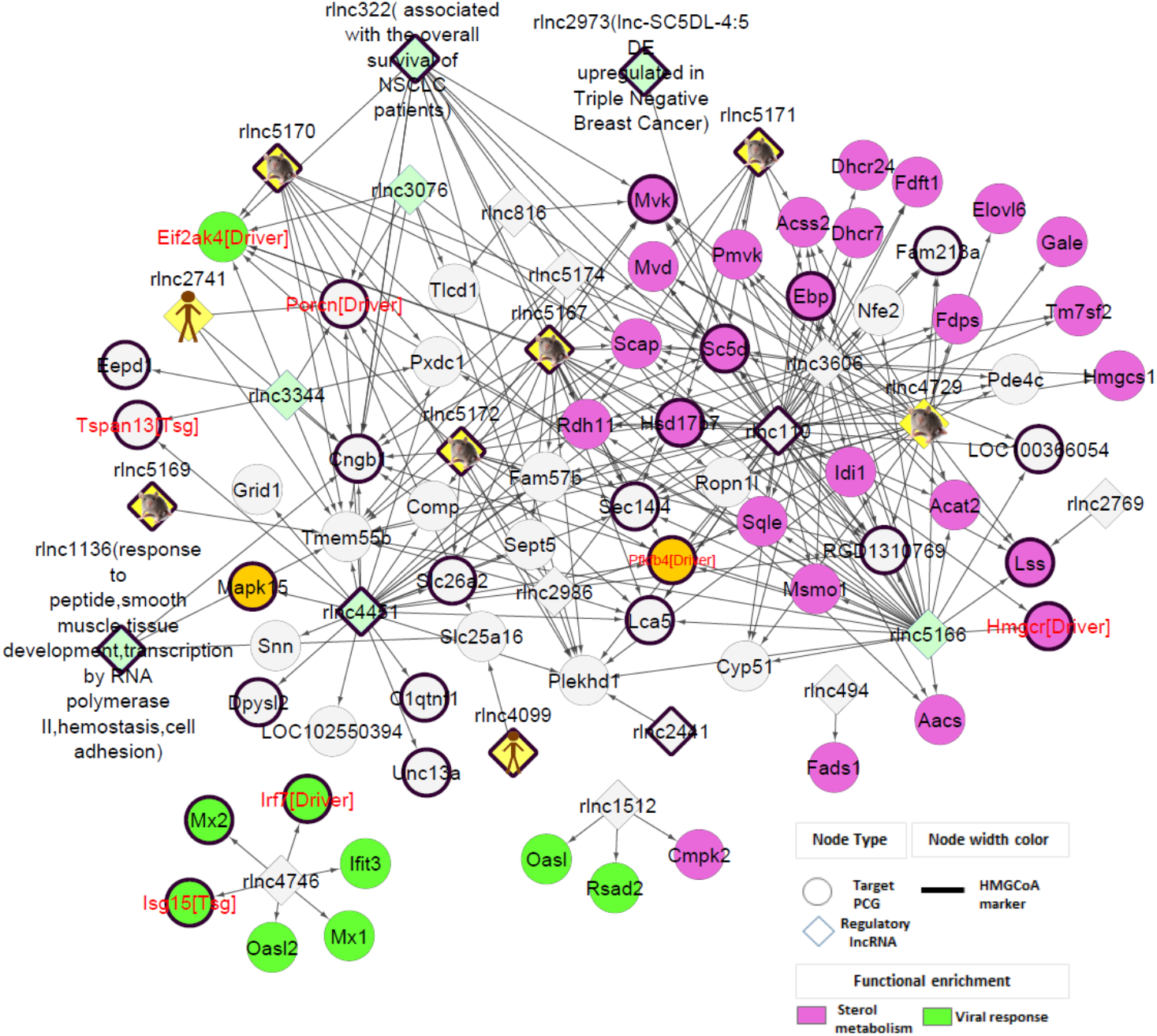
lncRNA−PCG causal network enriched for different biological processes. Each directed edge (arrows) represents a causal effect (absolute causal effect value > 0.5) of a xeno-lnc (diamond shapes) on the expression of a PCG. Ortholog information is represented by different node colors with node description added for functionally well−characterized lncRNAs.

### Xeno-lncs repressed by multiple xenobiotics may be hepatoprotective or hepatocarcinogenic

**–** Module C14 includes four of the five xeno-lncs consistently down-regulated by ≥ 20 xenobiotics (rlnc1425, rlnc715, rlnc2750, rlnc3088) (**Figure 8A-C**). Expression of these xeno-lncs correlates positively with several oncogenic drivers that are also widely down-regulated following xenobiotic exposure. The down-regulation of these xeno-lncs is thus a hepatoprotective response. For example, expression of rlnc1425, a module C14 hub gene that was down-regulated by 22 xenobiotics, showed positive correlations with oncogenic drivers Ptk6 and Ppard, both associated with hepatotoxicity [87-89] (**Figure 8A**). However, rlnc1425 also showed casual, negative correlation with metallothioneins Mt1a and Mt2a, which protect mice from hepatocarcinogen-induced liver damage and carcinogenesis [90] and are induced in rat liver, up to 16-fold, by the three ER agonists and by leflunomide (AhR) and econazole (CAR/PXR) (**Table S15A**). Expression of rlnc715, which was down-regulated by 21 chemicals, showed positive causal associations with cancer driver Ptk6 (<u>X. Chen et al. 2016</u>), and tumor suppressor gene Npas2; its impact on hepatoprotection vs. hepatotoxicity is thus uncertain (**Figure 8A, Table S15B**).

**Figure 8.**
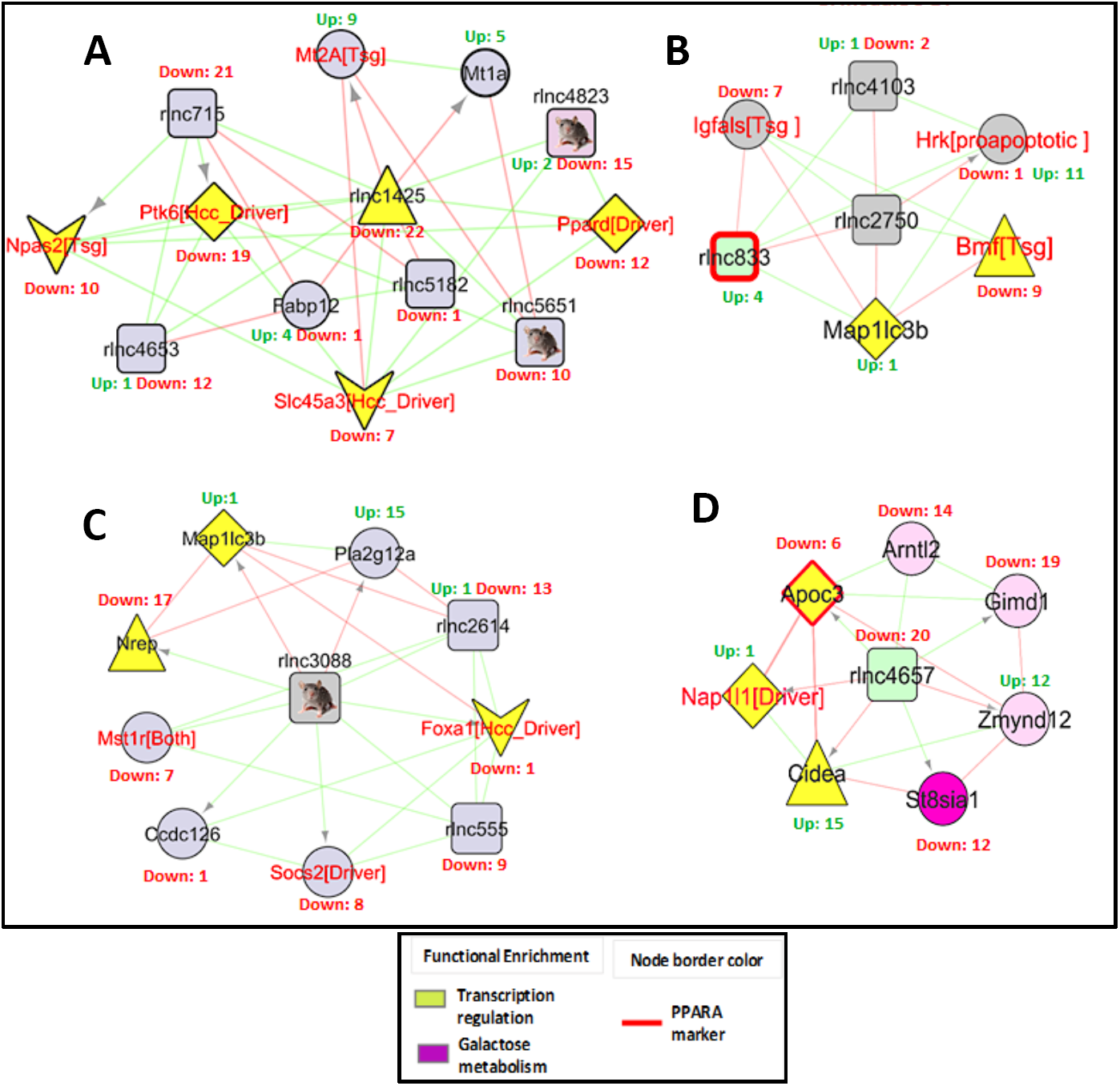
Sub-networks containing five xeno-lncs responsive to 20 or more multiple chemicals and their gene associations in network modules. Shown are sub−network derived from module C14 along with all direct connections and causal relationships (marked by directed edges) for**: A.** rlnc1425 and rlnc715, **B.** rlnc2750, and **C.** rlnc3088. **D.** Sub−network derived from module C12 showing all causal and correlation based associations for rlnc4657.

Expression of rlnc2750, which was down-regulated by 21 chemicals, was positively associated with the tumor suppressors Igfals [91] and Bmf [92] (**Figure 8B**), which were also down-regulated by multiple chemicals (**Table S15C**). Down-regulation of Igfals enhances IGF signaling and thereby promotes hepatocarcinogenesis [93], while down-regulation of Bmf is associated with hepatocellular carcinoma progression [94]. rlnc2750 had negative causal association with the pro-apoptotic gene Hrk, which was up-regulated by 11 xenobiotics and activates apoptosis under stress [95]. rlnc3088 was down-regulated by multiple chemicals, as were its causally associated target genes. These include the oncogenic driver Socs2 and the hub gene Nrep (P311) (**Figure 8C**), which plays a key role in reactive oxygen species-mediated hepatic stellate cell migration during liver fibrosis [96]. rlnc3088 had negative causal effects on Pla2g12a, a phospholipase A2, which was induced by 15 of the 27 chemicals, including the AhR agonist leflunomide (**Table S15D**).

The fifth lncRNA consistently down-regulated by >20 chemicals, rlnc4657, is in module C12, where it forms positive causal associations with three PCGs down-regulated by multiple chemicals (Apoc3, St8sia1, Gimd1) (**Figure 8D, Table S15E**). Apoc3 contributes to cardiovascular disease risk by increasing plasma triglycerides via lipolysis of triglyceride-rich lipoproteins [97], and its repression by PPAR activators is therapeutically beneficial. Gimd1, whose function is unknown, also showed strong, consistent repression by PPAR activators (log_2_ FC = −5 to −7) (**Table S3B**). Beneficial effects of rlnc4657 down-regulation by diverse chemicals are also apparent from the causally associated down-regulation of St8sia1, a ganglioside D3 synthase that promotes growth and metastasis in triple negative breast cancer [98]. St8sia1 was repressed by chemicals with various MOAs, including all three ER agonists, consistent with its repression by estradiol in human breast cancer cells [99]. The down-regulation of rlnc4657 by many xenobiotics may also have deleterious health effects. Thus, rlnc4657 showed negative causal association with two PCGs induced by multiple chemicals, one of which, Cidea, controls lipid storage, lipolysis and lipid secretion [100] and promotes hepatic steatosis when up-regulated in mouse liver [101]. The second gene, Zmynd12, is a zinc finger protein of unknown function. Finally, rlnc4657 was co-expressed with Arntl2 (Bmal2), which was repressed by 14 xenobiotics (**Figure 8D**). Arntl2 is an anti-apoptotic factor that is down-regulated in hepatocellular carcinoma, where its suppression enhances cell growth [102]. Thus, its repression by xenobiotics may contribute to hepatocarcinogenesis.

### Functional xeno-lnc candidates and predicted *cis* vs *trans* interactions with PCG targets

– We identified 67 top candidates for functional xeno-lncs (**Table S16A**) by integrating multiple datasets and criteria, including: their roles as causal regulators, or as hubs or bottlenecks in a functional module; shared orthology with well-characterized human or mouse lncRNAs; and xeno-responsiveness to multiple chemicals, or as a MOA-selective marker. Co-expression data and causal interactions between these 67 xeno-lncs and their putative PCG targets, combined with chromosomal location information, was used to predict whether each xeno-lnc regulates its targets in a *cis* or *trans* manner. Putative *cis* interactions were indicated for lncRNA−PCG pairs falling within a distance of 250 kb (**Table S16B**). *trans* interactions are shown **Table S16C**. The *cis* and *trans* PCG target lists were subdivided into activator and inhibitor effects, based on whether the xeno-lnc—PCG expression correlation was positive or negative. These analyses provide a basis for hypothesis-driven experimental studies on the functional roles of individual xeno-lncs and their downstream causal implications for responses to xenobiotic exposure.

## Discussion

LncRNAs regulate a wide range of cellular processes, including chromatin states, transcriptional output, mRNA stability and protein function [103]. However, the extent to which lncRNAs impact the toxicogenomic landscape is poorly understood. Little is known about their responses to xenobiotic exposure, and systematic approaches to deduce their regulatory roles in toxicological responses to foreign compounds have not been developed. Here we address these issues by introducing an integrated computational framework that utilizes transcriptomic data for discovery of global effects of xenobiotic exposure on pathways and mechanisms associated with dysregulation of lncRNA expression (**Figure 1**). We applied this framework to a rich toxicogenomic dataset comprised of 115 RNA-seq datasets representing 27 chemical exposures in a rat model [6] to assemble the toxico-transcriptome of liver, a key tissue for xenobiotic metabolism and toxification/detoxification. We discovered gene and isoform structures for almost 6,000 liver-expressed rat lncRNAs, a majority of which are novel genes, and established expression patterns for more than 1,400 xenobiotic-responsive lncRNAs (xeno-lncs), many with human and/or mouse orthologs. Further, we used two powerful data-driven approaches for co-expression analysis, WGCNA [31] and MEGENA [34], to discover xenobiotic-responsive gene modules enriched in various cellular processes, including fatty acid and sterol metabolism, cell cycle and immune response. Xeno-lncs occupying key positions as hubs or bottleneck genes were identified in the derived networks, and causal inference was used to identify xeno-lncs that causally influence expression of their target genes, i.e., are causal regulators of the biological network. Thus, we present a comprehensive toxicogenomic analysis of the effects of foreign chemicals on the non-coding transcriptome, and we elucidate xenobiotic-responsive regulatory lncRNAs for key biological pathways commonly perturbed in xenobiotic-exposed liver.

We used two complementary approaches for lncRNA discovery [9, 22] to increase the likelihood of discovering novel liver-expressed lncRNAs. 1,447 of the 5,795 rat liver-expressed lncRNAs identified were characterized as xeno-lncs based on their responsiveness to one or more xenobiotic exposures. We applied a relatively stringent threshold (>4-fold increase or decrease in expression at FDR <0.05) to reduce false positives resulting from transcriptional noise from lncRNAs expressed at a low level. The set of 1,447 xeno-lncs discovered here is defined by the transcriptional responses stimulated by the 27 chemical exposures included in our analyses, and should thus be viewed as a minimal xeno-lnc gene set. Additional xeno-lncs are very likely to be discovered and their gene structures and isoform models further refined once expression data for additional xenobiotic exposure datasets become available and can be integrated into the liver toxico-transcriptome presented here.

We found 81 xeno-lncs and 81 xenobiotic-responsive PCGs that were primarily associated with chemicals linked to a single MOA (xenobiotic MOA-selective marker genes; **Table S10**). We also found, however, that many xeno-lncs responded in common to multiple xenobiotics encompassing multiple MOAs. For example, 123 xeno-lncs each responded to at least 10 different chemicals (**Figure 2D)**, i.e., they respond via at least two different MOAs. While some of these xeno-lncs respond to chemicals that activate mechanistically related MOAs with overlapping target gene specificities (e.g., the nuclear receptors CAR/PXR and PPARA) [104, 105], many individual xeno-lncs respond to multiple chemicals that act via diverse MOAs. Such xeno-lnc responses are likely to encompass more general cellular and tissue responses to xenobiotic exposure, such as liver injury and liver repair, or the activation of hepatoprotective pathways and mechanisms. For example, four of the five xeno-lncs repressed by at least 20 of the 27 exposures, and encompassing all 7 MOAs, were positively co-expressed with several oncogenic drivers but showed either positive or negative associations with several tumor suppressors (**Figure 8**). Similar patterns of response to diverse xenobiotics were seen with some PCGs. Two PCGs active in xenobiotic detoxification, Sult2a2 and Ugt1a2, were up-regulated by ≥ 20 xenobiotics, while 22 of the 27 chemicals examined (including all MOAs except for DNA damage agents), strongly down-regulated carbonic anhydrase I (Car1), which like carbonic anhydrases III and VII, may protect liver from oxidative stress [106]. Leukotriene C4 synthase (Ltc4s) was also strongly repressed by 22 of the 27 chemicals tested. Ltc4s catalyzes biosynthesis of leukotriene C4, a potent inflammatory mediator [107], and its down-regulation protects from hepatic ischemia reperfusion injury [108]. Thus, xenobiotics that work through different MOAs can dysregulate common sets of lncRNAs and PCGs involved in xenobiotic detoxification and hepatoprotection.

Agonists of AhR, and the hepatotoxic chemical aflatoxin B1, which also has AhR agonist activity [56], often showed effects that were opposite of chemicals that act via other MOAs, most notably for chemicals that activate PPAR (**Figure 2C**). For example, the xenobiotic-metabolizing P450 enzyme *Cyp2b1* was induced by 5 of 6 PPAR agonists (**Table S3C**) but was repressed by all three AhR agonists, while *Acot1*, a PPAR target and a key player in rodent liver tumorigenesis [109], was up-regulated by 22 chemicals but not by the AhR agonists or by aflatoxin−B1. Similarly, we found five xeno-lncs that were each down-regulated by at least 20 of the 27 chemicals examined, but did not respond to the AhR agonists β-naphthoflavone and 3-methylcholanthrene. Mechanistic studies will be required to elucidate the mechanism for these disparate responses to activation of AhR vs. other MOAs.

In some cases, xeno-lnc orthologs showed significant species-specific responses to chemical exposures. Thus, many of the 140 orthologous pairs of rat and mouse lncRNAs that responded to CAR/PXR activators showed species-specific responses. This finding parallels the species difference in xenobiotic responses sometimes seen for orthologous PCGs (**Figure** S4), which may reflect factors such as species differences in the specificity of xenobiotic receptors for agonists and/or their target genes [30, 110-112]. This potential for species-dependent responses must be considered when evaluating lncRNA orthologs and their biological activities.

We built gene co-expression networks (gene modules) based on gene co-expression patterns for xeno-lncs and xenobiotic-responsive PCGs across the 27 xenobiotic exposures, and then used the associations between xeno-lncs and PCGs within the networks to infer the functions of liver-expressed xeno-lncs (guilt-by-association). Networks were reconstructed using WGCNA, which is widely used [10, 113-115], and using a complementary approach, MEGENA, which has identified mutational drivers in non-alcoholic fatty liver disease [116, 117]. Many of the gene modules we discovered were enriched in specific biological functions, including fatty acid metabolism, cell division, and immune response pathways. In some cases, MOA marker genes were grouped in a common module based on their coordinated co-regulation pattern (e.g., ER marker genes in module C10; **Figure S5**). We also implemented network modeling to identify hub or bottleneck genes that may regulate the overall network. Established cell cycle regulators (Bub1b, Prc1, Cdk1) occupied key positions in the cell division enriched module (**Figure** 5), and known regulators of fatty acid and lipid metabolism (Hadhb [118], Elovl6, Acot2, Acot1 [119], Pklr [116]) were top hubs in the network module enriched for this metabolic pathway (**Figure 4**). This validation of our approach lends support for the roles we propose for key xeno-lncs identified as hubs or bottlenecks in these biological processes and pathways. Key findings include the discovery of novel regulatory xeno-lncs that were co-expressed with tumor suppressors or cancer drivers in the immune response module C7 (**Figure S8)** or that occupied critical positions in networks enriched for fatty acid metabolism (**Figure 4**), viral response and sterol metabolism **(Figure 7).** In many cases, our findings were strengthened by using Parallel IDA, a causal inference algorithm [120-122], which can help distinguish true, causal regulatory relationships from simple correlative associations, as summarized in **Table S16A**.

Ortholog analysis, based on either sequence conservation or synteny [123], can facilitate discovery of biologically relevant properties of lncRNAs [124]. We used this approach to identify rat xeno-lncs showing conservation with known oncogenic lncRNAs, including H19 (rlnc449), LINC00665 (rlnc5324) [125], SNHG20 (rlnc 3767) [126], and Cytor (rlnc1439) [127] (**Table S4**). Chemicals that dysregulate these xeno-lncs can be expected to have a major impact on tumorigenesis. For example, rlnc1439/Cytor, whose up-regulation correlates with hepatocellular carcinoma progression and poor patient prognosis [127], was strongly induced by the CAR/PXR activators miconazole and methimazole and by chemicals that induce DNA damage (ifosfamide and N-nitrosodimethylamine) (**Table S14B**). Supporting this finding, the module C7 co-expression network containing rlnc1439 (**Figure S8B**) shows positive correlation between rlnc1439 and S100a11, which is overexpressed in many human cancers [128], and also with Sox4, which promotes hepatocarcinogenesis by inhibiting p53-mediated apoptosis [83].

In conclusion, we present a novel approach to discover xenobiotic-responsive lncRNAs and obtain important insights into their roles in diverse cellular processes, the biological pathways they impact and their effects on hepatic responses to xenobiotic exposures. The computational framework that we propose (**Figure 1**) can help prioritize lncRNA targets for further computational and experimental analysis, including toxicological risk assessment. These approaches are expected to advance the goals of computational toxicology, which requires new integrative approaches to measure, model and evaluate the toxicological consequences of xenobiotic exposure, including hazard potential and health risk assessment [129]. Future studies can apply this framework to discover regulatory lncRNAs in other contexts, including single cell analysis, which may be used to further characterize lncRNA responses and mechanisms of xenobiotic action across different cell types in liver, including zonated hepatocytes and endothelial cells [130], and may increase the reliability of co-expression and causal analysis by increasing the number of data points.

## Acknowledgements

This work is supported in part by NIH grant ES 024421 (to DJW).

## Figure legends

**Scheme 1.**
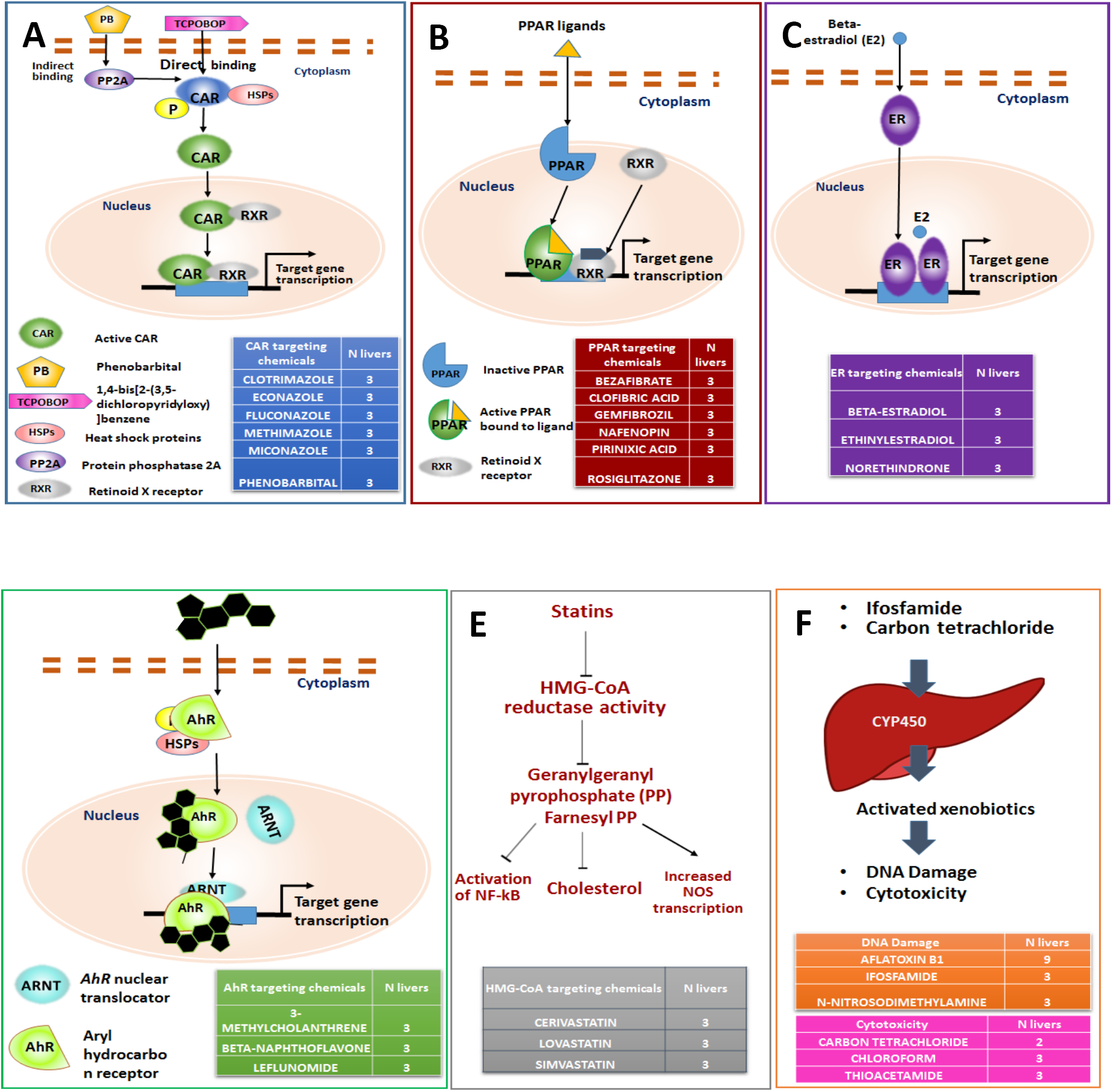
Modes of actions (MoA) for 27 xenobiotic exposures. Shown are schematic diagrams of thre MOA of each set of chemicals, and the numbers of livers included in each dataset. **A**, CAR/PXR; **B**, PPAR; **C**, ER; **D**, AhR; **E**, HMG-CoAR; **F**, cytotoxoicity and DNA damage agents.

## Supplemental figures

**Figure S1.**
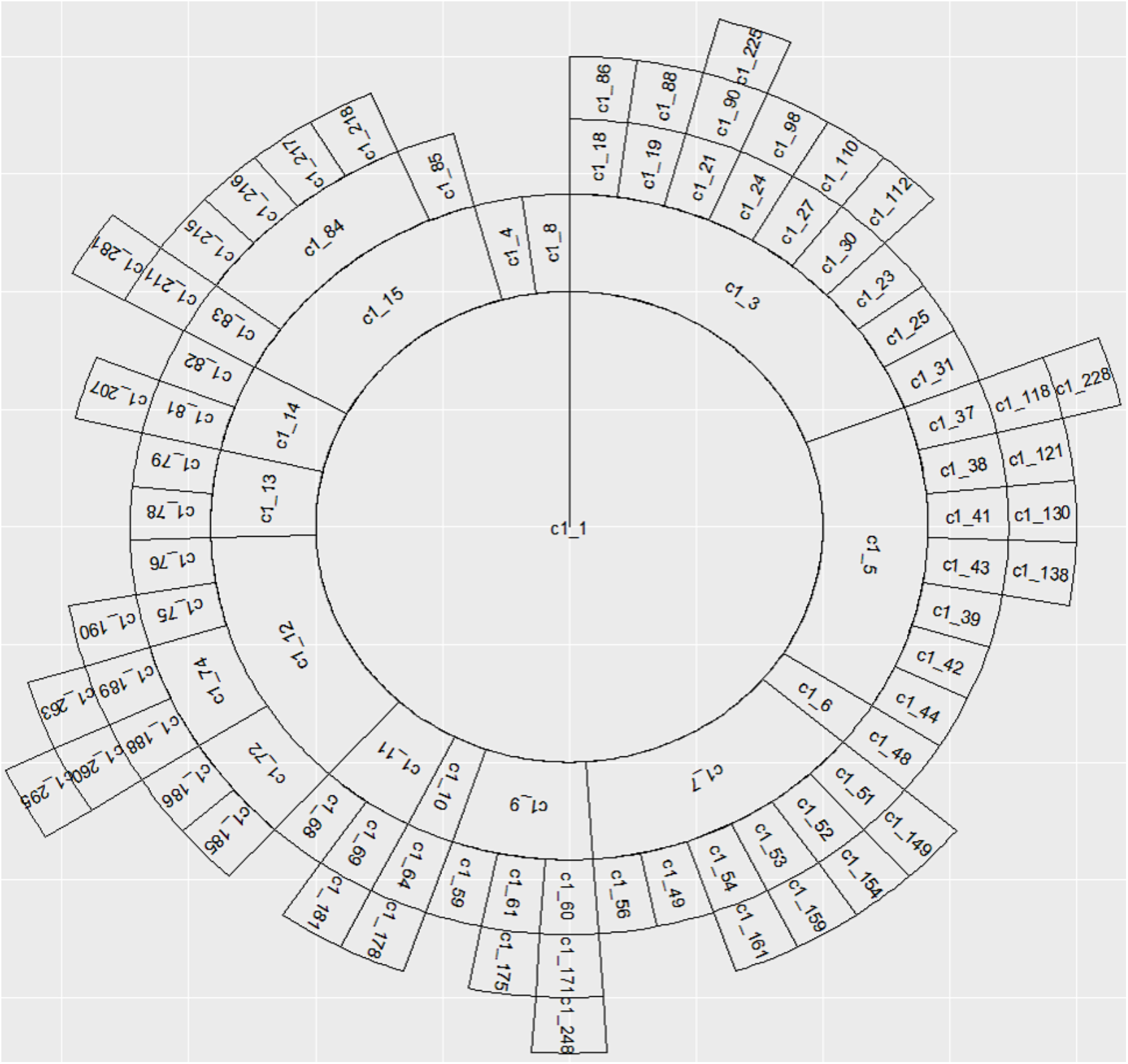
Module and sub−module hierarchy for MEGENA modules. MEGENA uses a divisive clustering approach and discovers co-expression modules in a multi-layer manner. The innermost core, C1_1, contains all 2,637 PCGs and 1,447 xeno-lncs, which are clustered into 13 gene modules in layer 1 (C1_3 to C1_15). Gene modules in layer 1 are further clustered into smaller compact sub−modules in layer 2. This process continues until no further compact child clusters are formed.

**Figure S2.**
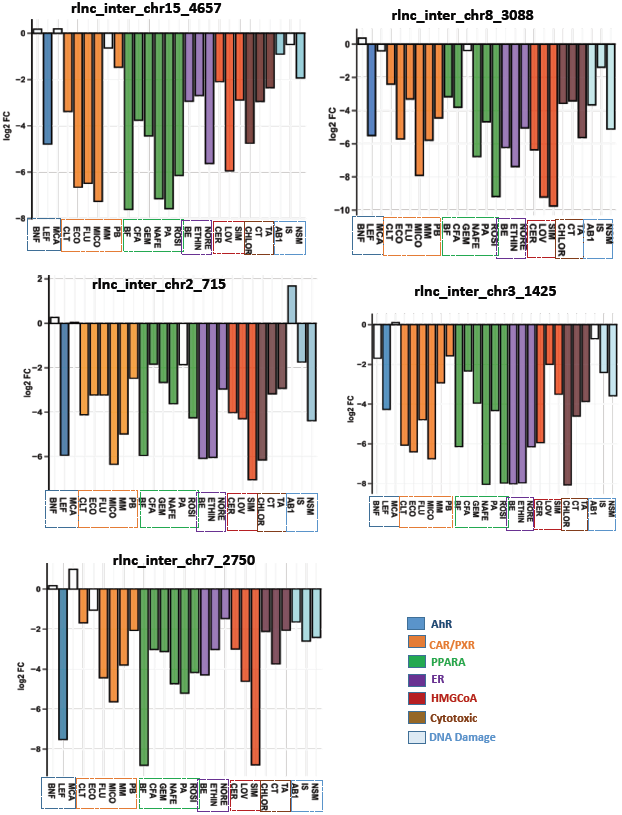
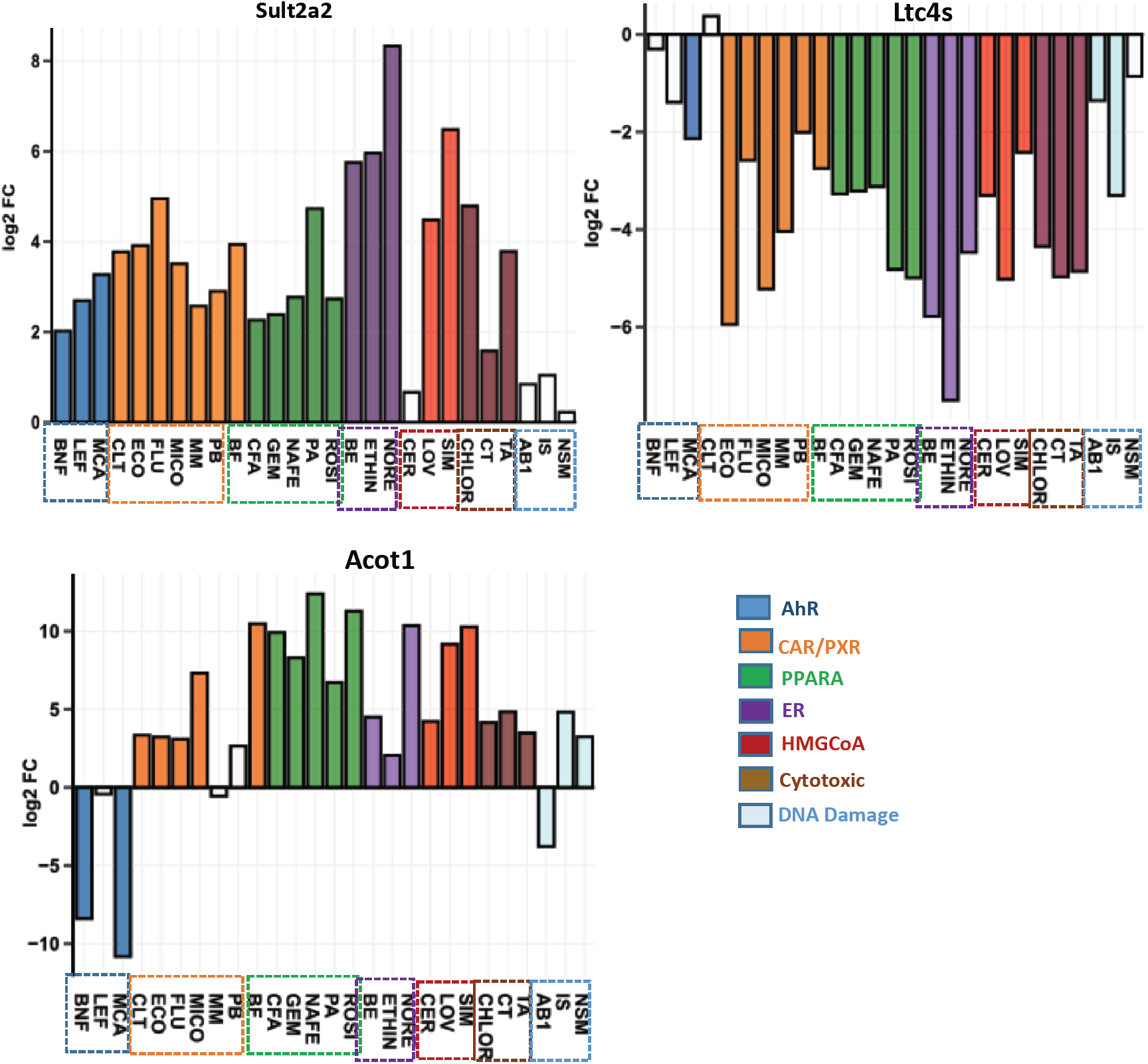
Gene expression data for xeno-lncs (A) and protein coding genes (PCGs) (B) that are consistently induced or repressed by ≥20 of the 27 chemicals examined. Data are shown as log2 fold-change (FC) values along the Y-axis. Bars shown in white, FDR < 0.05. **A.** Five lncRNAs (rlnc4657, rlnc3088, rlnc715, rlnc1425, and rlnc2750) showed down regulation in 20 or more chemicals. **B.** Sult2a was up regulated by 22 out of 27 chemicals, and Ltc4s gene was down-regulated by 22 out of 27 chemicals. Acot1 was up-regulated by 21 chemicals, but was down-regulated by two of the three AhR agonists, and by aflatoxin−B1, which also has AhR agonist activity (see text). Bars are colored according to the MOA of each chemical.

**Figure S3.**
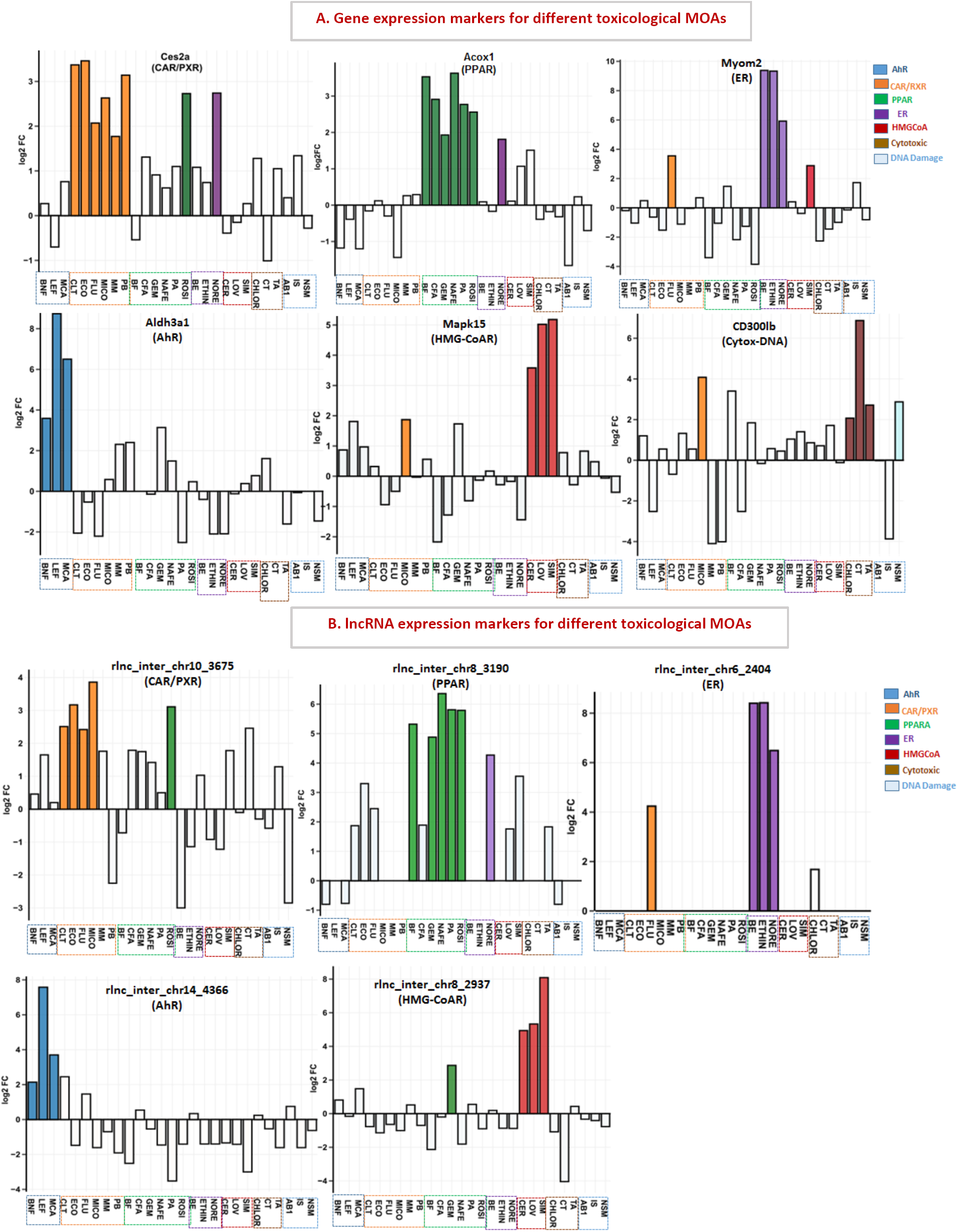
Gene expression profiles across all 27 chemicals for representative MOA-selective marker genes, including PCGs (A) and xeno-lncs (B). Gene expression data is in log_2_ FC values (y-axis), and bars are colored according to the MOA of each chemical. White bars, FDR < 0.05.

**Figure S4A.**
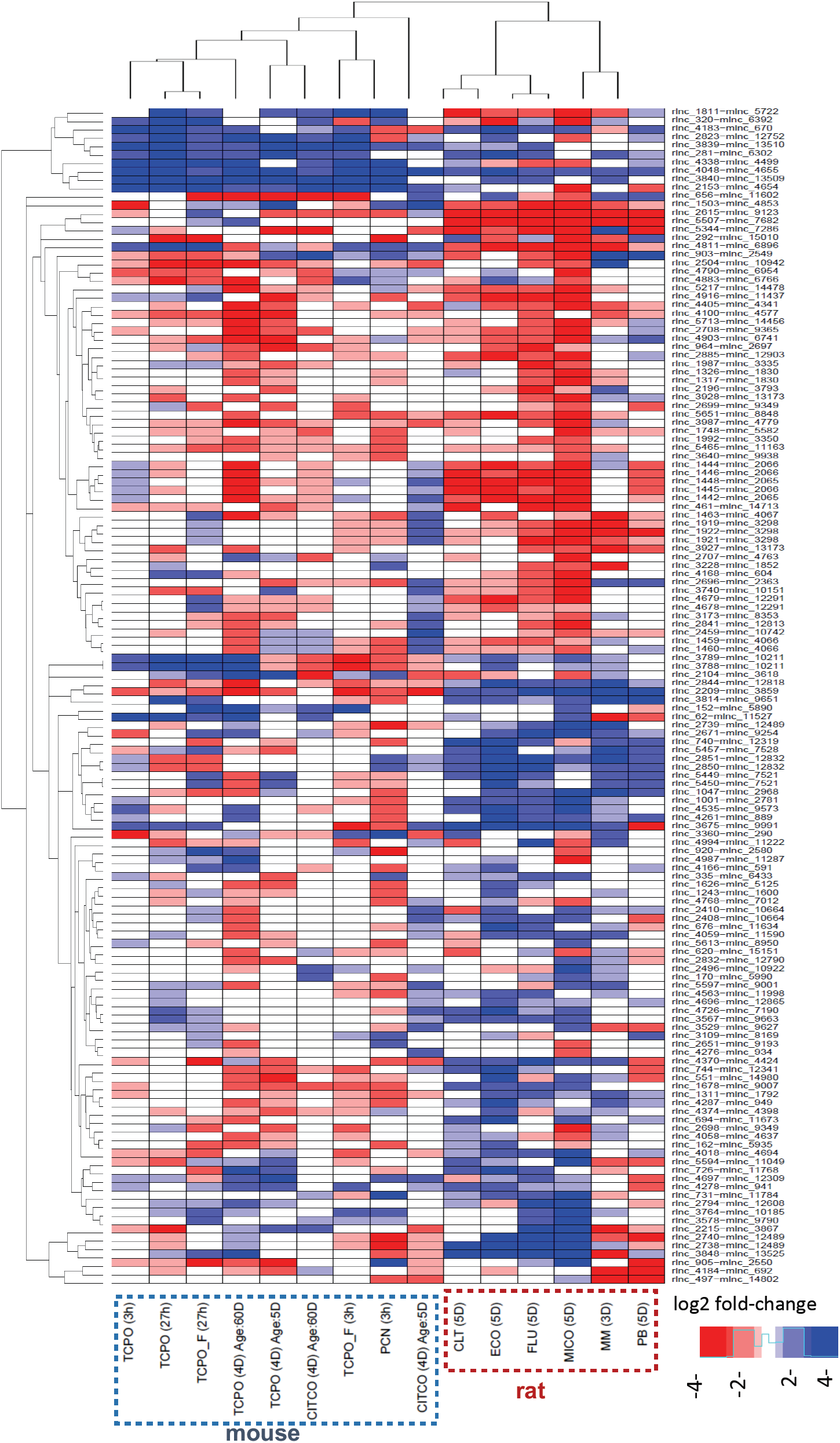
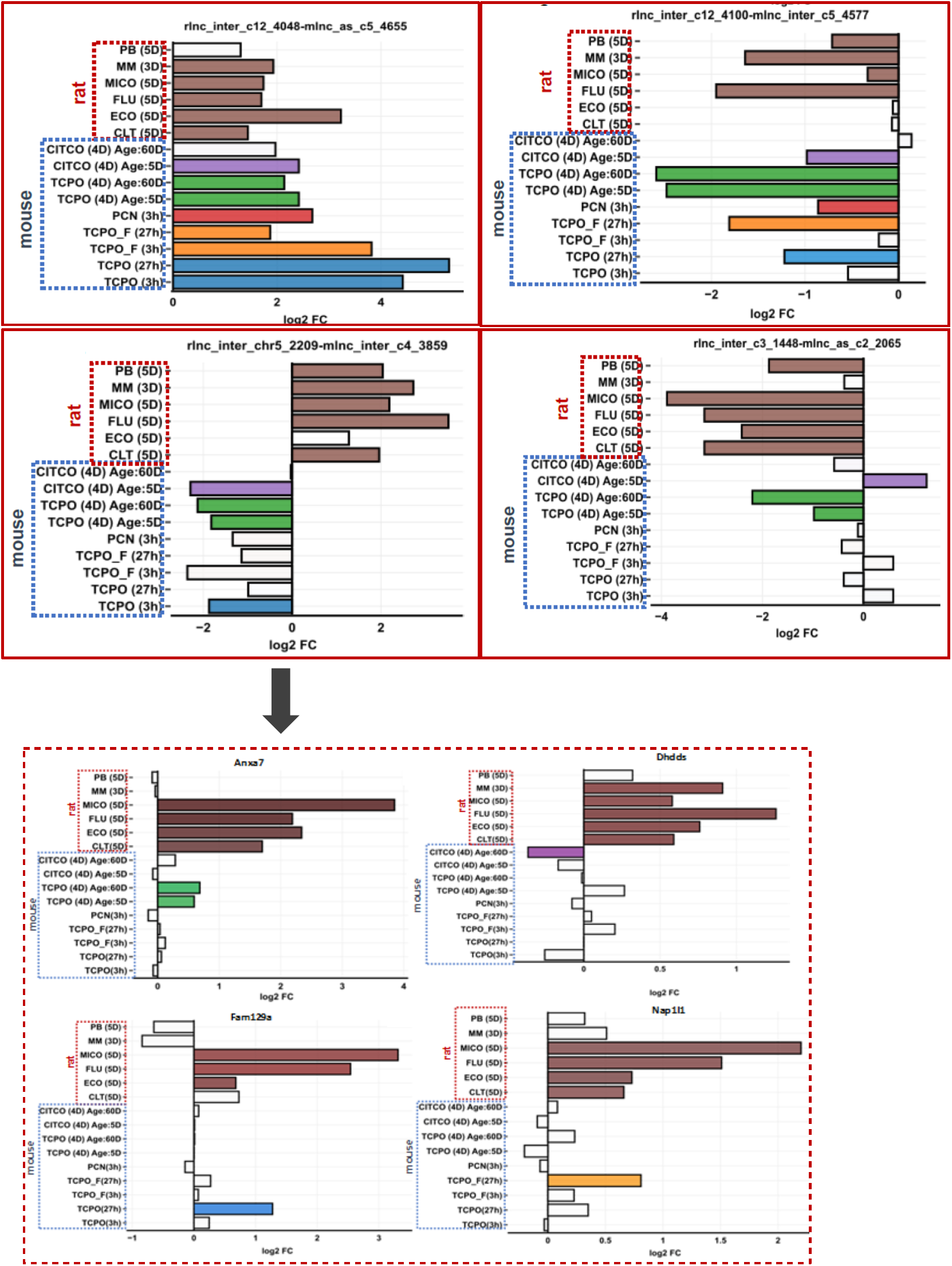
Rat−mouse ortholog responses to xenobiotics that are activators of CAR or PXR. **A**, Heatmap of 140 rat xeno-lncs whose mouse orthologs was significantly dysregulated by a CAR or PXR agonists in one of the mouse datasets (see text). Data are displayed by hierarchical clustering using Euclidean distance metric and Ward.d2 minimum variance criterion. Each row represents a lncRNA rat−mouse ortholog pair and each column represents one gene expression dataset. **B**, Expression data for select rat-mouse xeno-lnc orthog pairs (top) and of four PCGs co-expressed with the orthologs pair rlnc2209-mlnc3859 (bottom).

**Figure S5.**
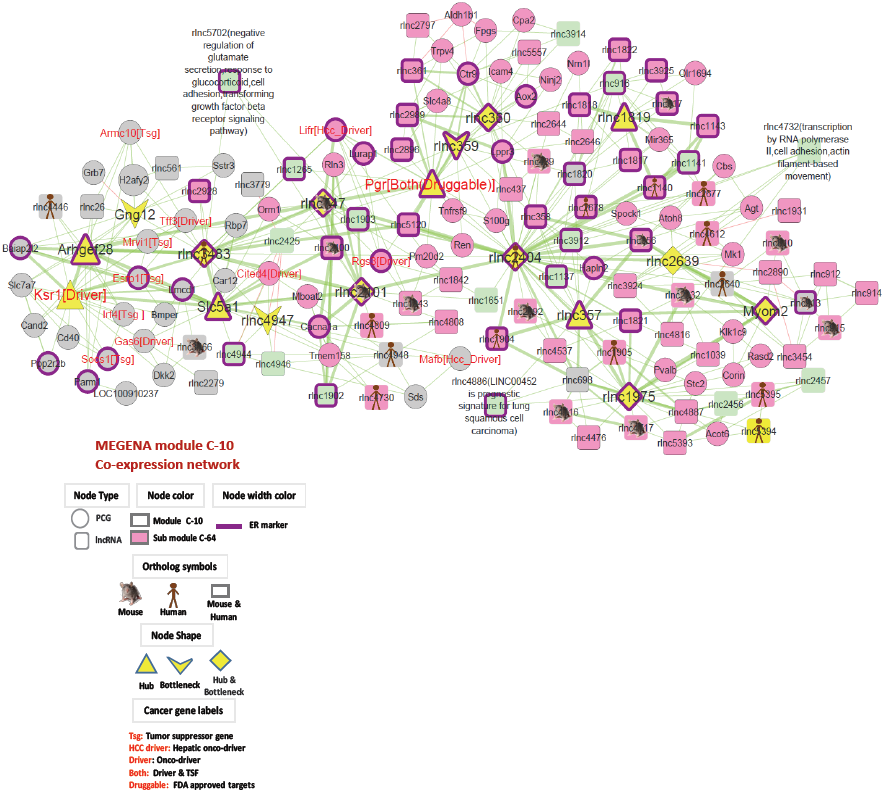
Module C10, which is highly enriched in ER marker genes. Xeno-lncs occupying central position as hubs and bottlenecks for module C10 that contained all 50 ER markers.

**Figure S6.**
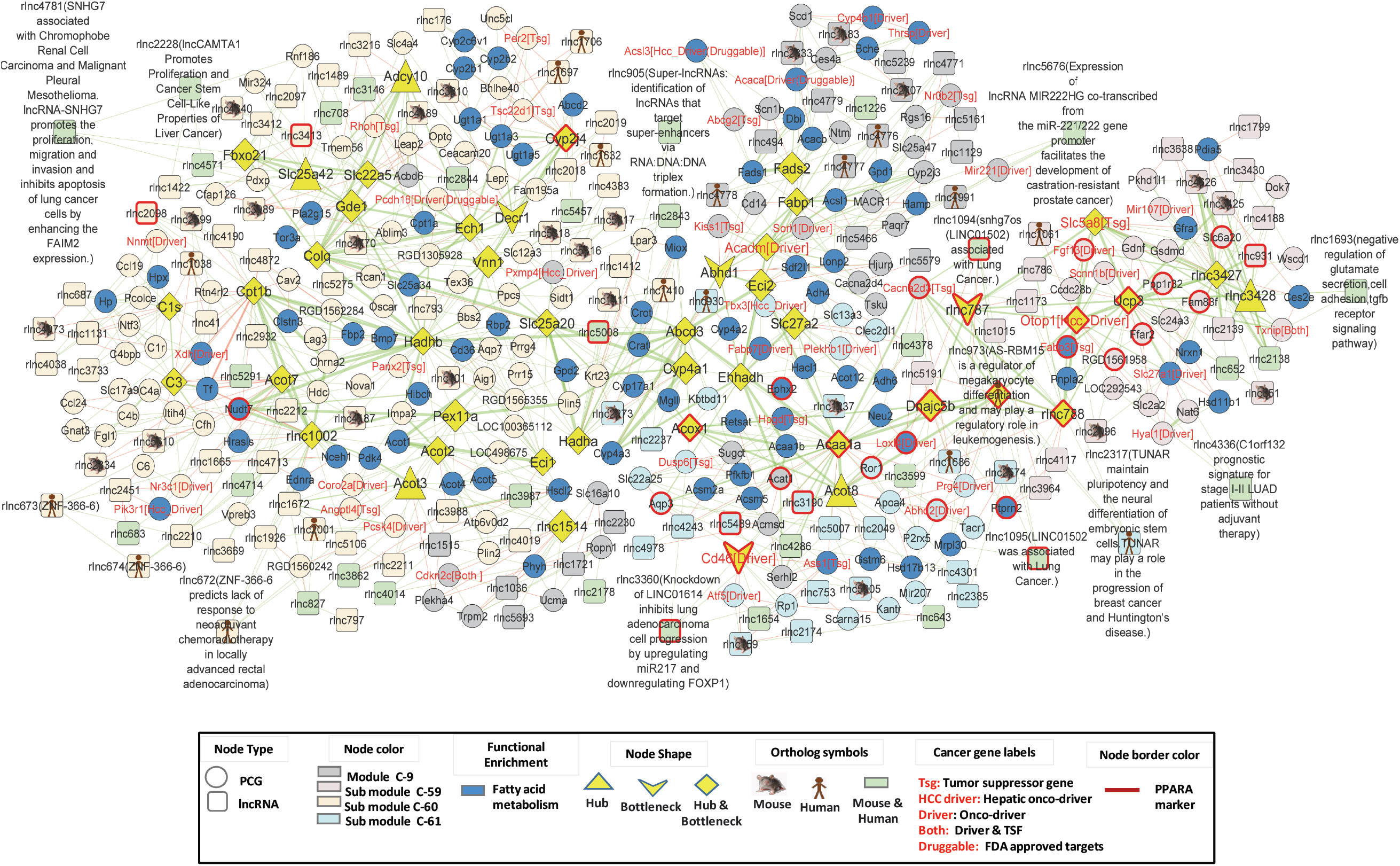
Complete network of module C9, which is enriched for fatty acid metabolism terms. Fatty acid metabolism genes are represented by nodes shown in blue. Network submodules (C−59, C−60, C−61) are represented by different colors.

**Figure S7.**
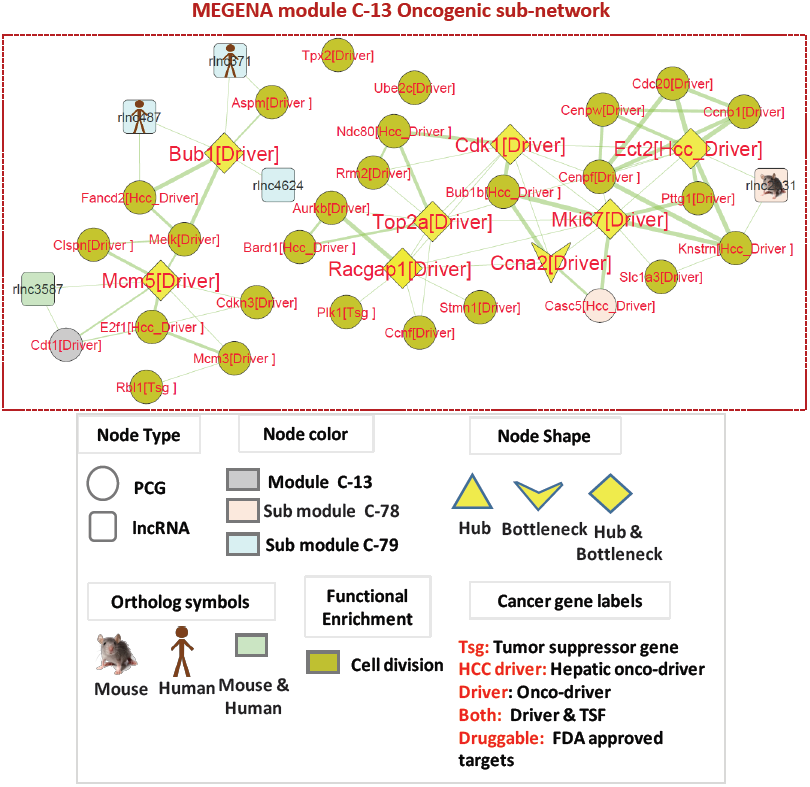
Oncogenic sub−network derived from module C13. This module includes 35 PCGs with cancer−associated roles, either as oncogenic drivers or tumor suppressors (TSGs). Five xeno-lncs are connected directly to these genes.

**Figure S8A.**
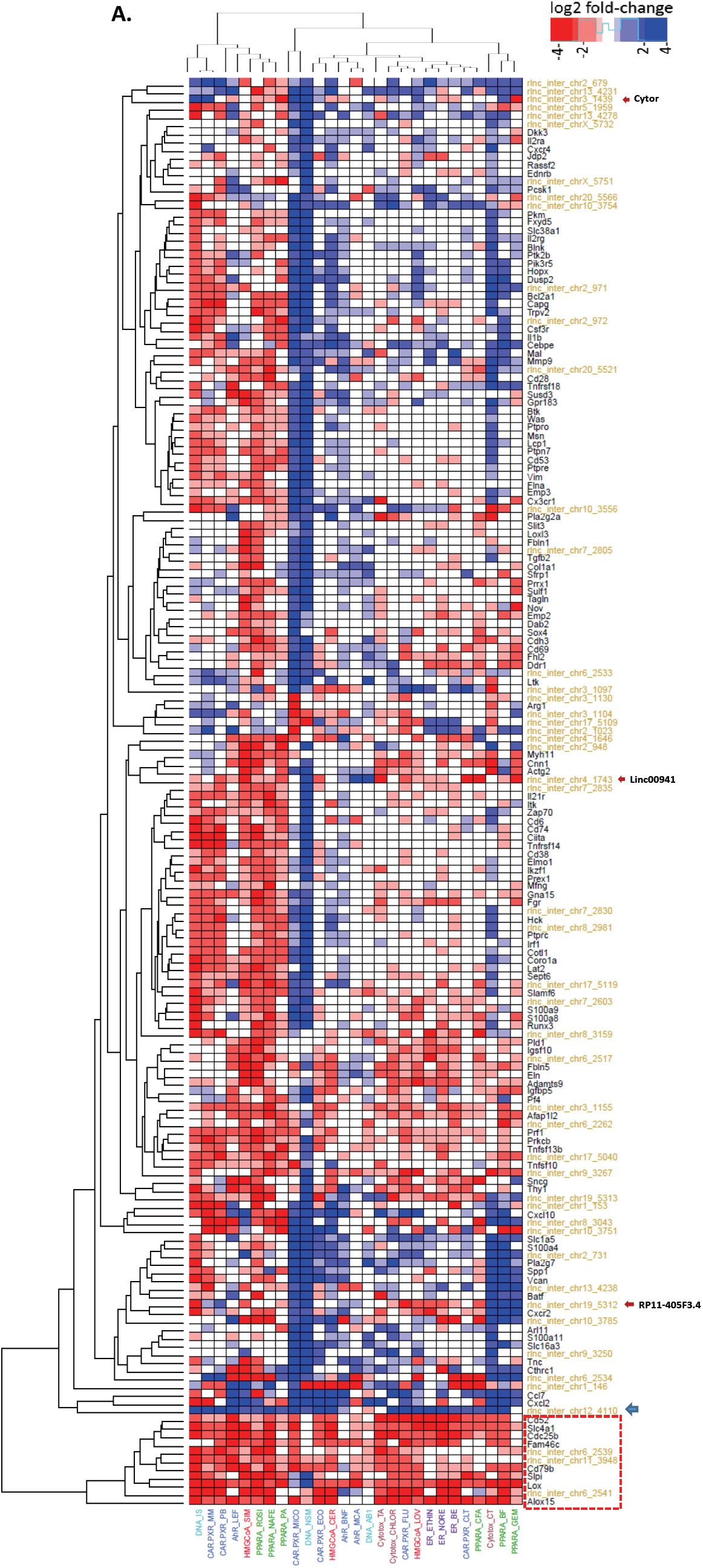
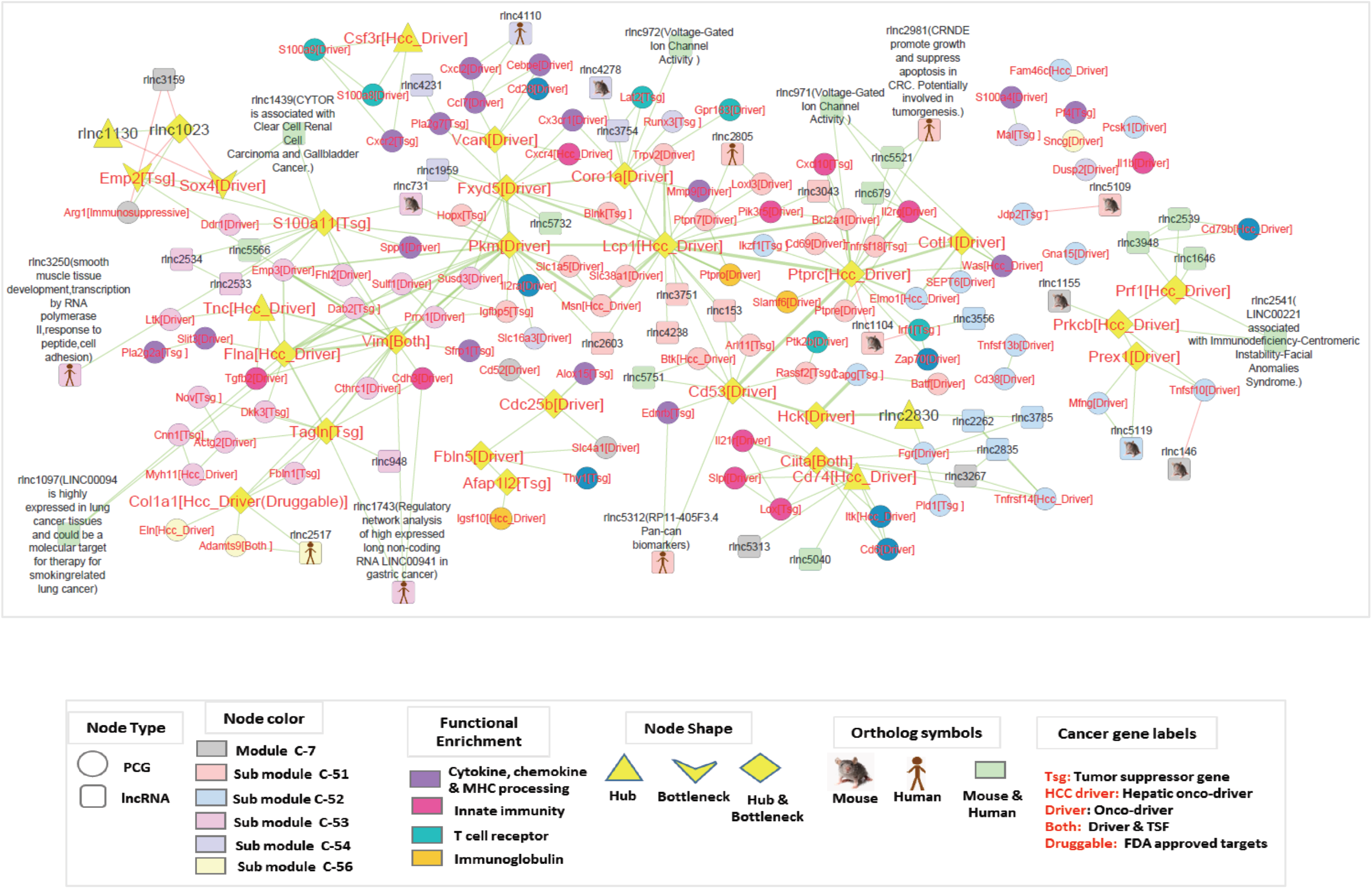
Heatmap presenting gene response for oncogenic genes from Module C7, which is enriched for immune response genes. The 125 oncogenes (black text) genes displayed here are connected to one or more of 49 xeno-lncs (gold text). We observed a small cluster (marked at the bottom as a dotted rectangle) that was robustly down-regulated across all chemical exposures. Xeno-lnc rlnc4110 (blue arrow) was induced across all conditions. In addition, we identified several known lncRNA orthologs (red arrows). **B**. Oncogenic gene sub-network, excerpted from module C7. This network presents oncogenic genes and their direct xeno-lnc neighbors. Three of the onco-lnc orthologs shown, lnc-CYTOR, Linc00941, RP11-405F3.4, are connected to critical node genes in the network.

